# PARP16 protects against pathological cardiac remodelling by ADP-ribosylation-dependent inhibition of NFAT transcription factor

**DOI:** 10.64898/2026.03.30.715447

**Authors:** Sima Zarinfard, Sukanya Raghu, Arathi Bangalore Prabhashankar, Abhishek Chowdhury, Pavithra Jayadevan, Raksha Rajagopal, Anurag Sharma, Amarjeet Shrama, Preethi Amardeep Ray, Prasanna Simha Mohan Rao, Utpal Nath, Kumaravel Somasundaram, Michael Hottiger, Nagalingam R. Sundaresan

**Author notes:** **Correspondence should be addressed to:** Nagalingam Ravi Sundaresan: SB-02, Department of Microbiology and Cell Biology, Biological Sciences Building, Indian Institute of Science, Bangalore, Email ID.

## Abstract

Mono-ADP-ribosylation is a post-translational modification that regulates diverse cellular processes. PARP16 is an endoplasmic reticulum-associated mono-ADP-ribosyltransferase implicated in stress-response signalling; however, its role in cardiac remodelling and dysfunction has not been fully defined. Our results suggest that PARP16 expression was reduced in human heart failure samples. Deletion of PARP16 in several mice models promoted ventricular dilatation, fibrosis, fetal gene reactivation, and systolic dysfunction, whereas cardiomyocyte-specific overexpression of PARP16 attenuated isoproterenol-induced remodelling in mice. Transcriptomic profiling and functional studies identified NFAT signalling as a major downstream pathway activated following PARP16 deficiency. Specifically, loss of PARP16 increased nuclear accumulation, promoter occupancy, and transcriptional activity of NFAT1. Mechanistically, PARP16 interacted with NFAT1 and suppressed NFAT-dependent transcription through a catalytic activity-dependent mechanism. Proteomic, structural, and functional analyses identified E398 and T533 residues of NFAT1 associated with PARP16-mediated regulation. Furthermore, pharmacological inhibition of NFAT improved cardiac function and remodelling in PARP16-deficient mice. These findings indicate that PARP16 acts as a negative regulator of NFAT signalling and contributes to protection against adverse cardiac remodelling.

## Introduction

Heart failure remains a major global health burden and is frequently preceded by pathological cardiac remodelling, a process characterised by persistent alterations in cardiac structure, function, and gene expression that ultimately drive progressive ventricular dysfunction [1–7]. The transition from adaptive cardiac responses to maladaptive remodelling is governed by complex signalling networks and transcriptional programs that regulate cardiomyocyte growth, fibrosis, inflammation, and contractile function [8, 9]. Post-translational modifications are central to these regulatory circuits, and disruption of PTM-dependent signalling has emerged as a recurrent feature of cardiovascular disease[9]. Several PTMs, including SUMOylation [10], O-GlcNAcylation[11], S-nitrosylation[12], phosphorylation, and ubiquitination[9, 13], are now recognised as important determinants of cardiac homeostasis and remodelling.

ADP-ribosylation has gained prominence as a regulatory PTM in cardiovascular biology[14–16]. ADP-ribosyltransferases catalyse the transfer of ADP-ribose from NAD⁺ to target proteins, thereby influencing a broad range of cellular processes. PARP1, PARP2, and tankyrases primarily mediate poly-ADP-ribosylation[17], whereas several other PARP family members catalyse mono-ADP-ribosylation[18]. Prior studies have established important roles for PARP1 in cardiac remodelling and heart failure progression [14, 19, 20], and PARP2 has been implicated in cardiomyocyte ageing and remodelling [21, 22]. By contrast, the contribution of mono-ADP-ribosyltransferases to cardiac homeostasis and disease remains less well defined.

PARP16 is an endoplasmic reticulum-associated mono-ADP-ribosyltransferase that regulates PERK- and IRE1α-dependent unfolded protein response signalling[23]. Beyond its role in endoplasmic reticulum stress responses, PARP16 has been linked to vascular ageing and neointimal hyperplasia [24, 25]. More recently, PARP16 has been proposed to participate in pathological cardiac remodelling; however, the precise role of PARP16 in the heart remains incompletely defined. Although recent studies have linked PARP16 to cardiac stress responses [26], differences in the reported molecular mechanisms and disease-associated signalling pathways indicate that its function during cardiac remodelling is not yet fully resolved. These observations underscore the need for complementary genetic and mechanistic studies to define the physiological role of PARP16 in maintaining cardiac homeostasis.

Among the transcriptional pathways implicated in pathological cardiac remodelling, calcineurin-NFAT signalling is a key driver of maladaptive gene expression and cardiac dysfunction**[27].** NFAT activity is regulated by multiple post-translational modifications that control its localisation and transcriptional output **[28–32].** Previous studies have shown that ADP-ribosylation can modulate NFAT activity and DNA binding, raising the possibility that ADP-ribosyltransferases directly shape NFAT-dependent transcriptional responses during cardiac stress. However, the role of mono-ADP-ribosyltransferases in NFAT regulation remains unknown.

In this study, we investigated the role of PARP16 in pathological cardiac remodelling using whole-body, cardiomyocyte-specific, and inducible cardiomyocyte-specific PARP16 knockout mice, together with cardiomyocyte-specific PARP16 transgenic mice. We show that PARP16 loss promotes adverse cardiac remodelling and dysfunction, whereas PARP16 overexpression attenuates stress-induced remodelling. Mechanistically, we identify NFAT1 as a major downstream effector of PARP16 signalling and provide evidence that PARP16 suppresses NFAT1 transcriptional activity through a catalytic activity-dependent mechanism. Together, these findings define a PARP16-NFAT1 signalling axis in pathological cardiac remodelling.

## Results

### PARP16 expression is reduced in failing human hearts and attenuates hypertropy in cardiomyocytes

To assess whether PARP16 is associated with human heart failure and pathological cardiac remodelling, we examined its expression in human cardiac samples. Analysis of a publicly available transcriptomic dataset showed reduced PARP16 expression in interventricular septal tissue from patients with dilated cardiomyopathy compared with non-failing controls (Fig. 1A). In line with this finding, PARP16 mRNA and protein levels were significantly lower in failing human heart samples (Fig. 1B–D), indicating that PARP16 expression is reduced in human heart failure. To determine the functional consequences of PARP16 loss, we depleted PARP16 in neonatal rat cardiomyocytes (NRCMs). PARP16 knockdown increased perinuclear ANP localisation, disrupted α-actinin organisation, and increased expression of the fetal gene markers BNP and βMHC relative to control cells (Fig. 1E–G), supporting a remodelling-associated phenotype following PARP16 depletion.We next asked whether increased PARP16 expression alters cardiomyocyte responses to hypertrophic stimulation. Under basal conditions, PARP16 overexpression did not affect cardiomyocyte morphology or fetal gene expression (Fig. 1H–K). However, phenylephrine induced remodelling-associated changes in control cardiomyocytes were markedly attenuated in PARP16-overexpressing cells (Fig. 1I–K). Together, these findings indicate that PARP16 expression is reduced in failing human hearts and that PARP16 limits remodelling-associated responses in cardiomyocytes.

**Figure 1.**
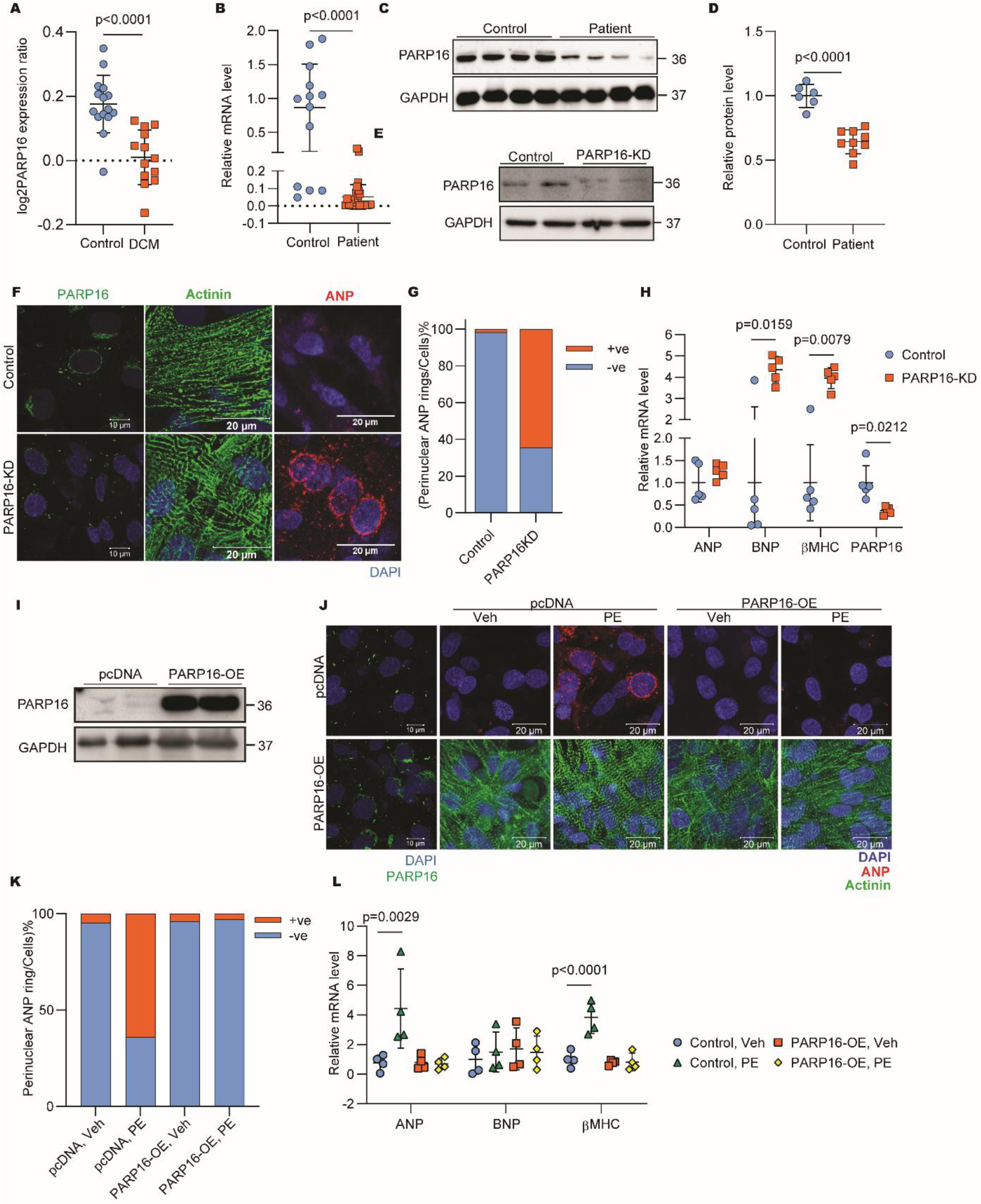
PARP16 is reduced in failing human hearts and restrains cardiomyocyte remodelling. (A) PARP16 mRNA expression in interventricular septum samples from non-failing controls and patients with dilated cardiomyopathy (DCM), analysed using a publicly available GEO dataset (GSE278958; n = 15 control and n = 13 DCM samples). (B) Quantitative RT-PCR analysis of PARP16 mRNA expression in left ventricular tissue from non-failing control and failing human hearts. Expression was normalised to GAPDH (n = 20 for controls and n = 10 for failing hearts). (C) Representative immunoblot of PARP16 protein expression in left ventricular tissue from non-failing control and pre-surgery failing human hearts. Molecular weight markers are indicated. (D) Densitometric quantification of PARP16 protein levels in a human Heart failure patient, normalised to GAPDH; n=6, 9. (E) Representative immunoblot showing PARP16 protein expression in neonatal rat cardiomyocytes (NRCMs) transfected with control siRNA or PARP16-targeting siRNA. (F) Representative confocal images of NRCMs transfected with control siRNA or PARP16-targeting siRNA and immunostained for atrial natriuretic peptide (ANP), α-actinin, and DAPI. Scale bar, as indicated. (G) Quantification of NRCMs exhibiting perinuclear ANP localisation under control and PARP16 knockdown conditions. (H) Quantitative RT-PCR analysis of ANP, BNP, and βMHC mRNA expression in control and PARP16-knockdown NRCMs (n = 5 independent experiments). (I) Representative immunoblot showing PARP16 protein expression in NRCMs transduced with either a control or a PARP16 overexpression construct. (J) Representative confocal images of control and PARP16-overexpressing NRCMs treated with vehicle or phenylephrine (PE) and immunostained for ANP, α-actinin, and DAPI. Scale bar, as indicated. (K) Quantification of NRCMs exhibiting perinuclear ANP localisation in control and PARP16-overexpressing NRCMs following vehicle or PE treatment. (L) Quantitative RT-PCR analysis of ANP, BNP, and βMHC mRNA expression in control and PARP16-overexpressing NRCMs treated with vehicle or PE (n = 5 independent experiments). Data are presented as mean ± SD. Statistical analyses were performed using unpaired two-tailed t-tests with Welch’s correction, Mann–Whitney tests, or two-way ANOVA followed by Sidak’s multiple-comparisons test, as indicated. Exact P values are provided in the source data. Source data are provided as a Source Data file.

### Whole-body deletion of PARP16 promotes pathological cardiac remodelling

To investigate the role of PARP16 in cardiac function in vivo, we generated whole-body PARP16 knockout (KO) mice (Fig. 2A, B). PARP16 KO mice showed increased heart size, as reflected by elevated heart weight-to-body weight (HW/BW) and heart weight-to-tibia length (HW/TL) ratios compared with control littermates (Fig. 2C, D). Echocardiographic analysis revealed increased left ventricular internal diameter (LVID) and reduced fractional shortening, whereas left ventricular posterior wall thickness (LVPW) was not significantly altered, consistent with adverse ventricular remodelling (Fig. 2E–G). Histological analysis showed increased cardiomyocyte cross-sectional area and interstitial fibrosis in PARP16 KO hearts compared with controls (Fig. 2H–J). In line with these structural changes, expression of the fetal gene markers BNP and βMHC increased in PARP16-deficient hearts (Fig. 2K). Together, these findings indicate that loss of PARP16 promotes pathological cardiac remodelling, characterised by ventricular dilatation, fibrosis, fetal gene reactivation, and impaired cardiac function.

**Figure 2.**
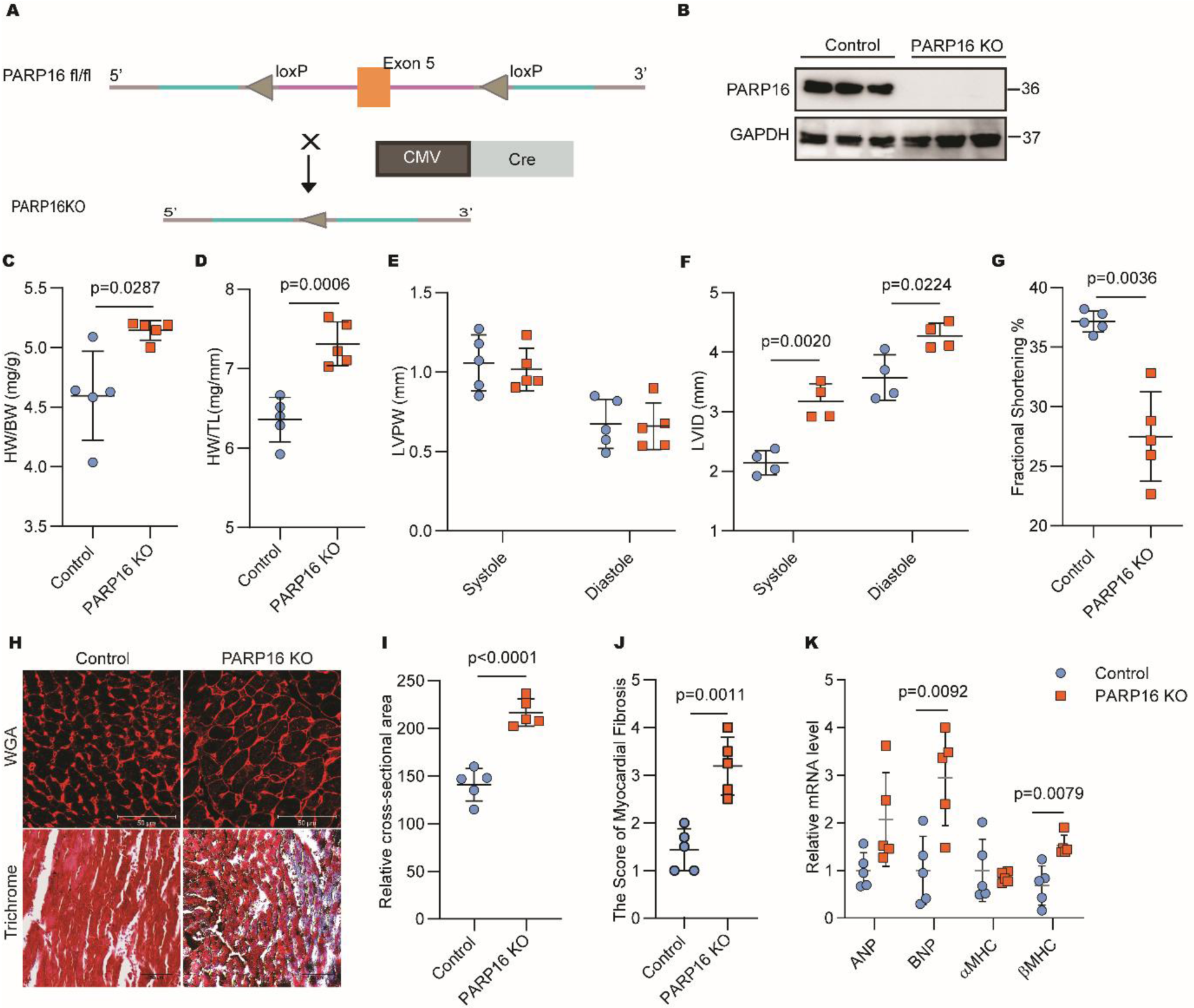
Whole-body PARP16 deficiency induces pathological cardiac remodelling. (A) Schematic illustrating the generation of whole-body PARP16 knockout (KO) mice by crossing PARP16 floxed mice with CMV-Cre mice. (B) Representative immunoblot of PARP16 protein expression in heart tissue from control and PARP16 KO mice. Molecular weight markers are indicated. (C) Heart weight-to-body weight (HW/BW) ratio in 5-month-old control and PARP16 KO mice. (D) Heart weight-to-tibia length (HW/TL) ratio in 5-month-old control and PARP16 KO mice. (E) Left ventricular posterior wall thickness (LVPW) at systole and diastole measured by echocardiography in 5-month-old control and PARP16 KO mice. (F) Left ventricular internal diameter (LVID) at systole and diastole measured by echocardiography in 5-month-old control and PARP16 KO mice. (G) Fractional shortening (FS) determined by echocardiography in 5-month-old control and PARP16 KO mice. (H) Representative wheat germ agglutinin (WGA)-stained heart sections for assessment of cardiomyocyte cross-sectional area and Masson’s trichrome-stained sections for assessment of cardiac fibrosis in hearts from 5-month-old control and PARP16 KO mice. Scale bar, 50 μm. (I) Quantification of cardiomyocyte relative cross-sectional area from WGA-stained heart sections. (J) Quantification of fibrotic score from Masson’s trichrome-stained heart sections. (K) Quantitative RT-PCR analysis of ANP, BNP, αMHC, and βMHC mRNA expression in hearts from control and PARP16 KO mice. Data are presented as mean ± SD. Statistical significance was determined using unpaired two-tailed t-tests with Welch’s correction for panels C–I and K (ANP, BNP, and αMHC) and the Mann–Whitney test for βMHC expression. n = 5 mice per group for all quantitative analyses unless otherwise indicated. Exact P values are provided in the source data. Source data are provided as a Source Data file.

### Cardiomyocyte-specific deletion of PARP16 recapitulates pathological cardiac remodelling

To determine whether the remodelling phenotype observed in whole-body PARP16 knockout mice was attributable to loss of PARP16 in cardiomyocytes, we generated cardiomyocyte-specific PARP16 knockout (csKO) mice (Fig. 3A–C). As in the whole-body knockout model, csKO mice exhibited increased heart size, as reflected by elevated heart weight-to-body weight (HW/BW) and heart weight-to-tibia length (HW/TL) ratios compared with control littermates (Fig. 3D–F). Echocardiographic analysis showed reduced left ventricular posterior wall thickness (LVPW), increased left ventricular internal diameter (LVID), and decreased fractional shortening in csKO mice, corroborating adverse ventricular remodelling and impaired cardiac function (Fig. 3G–I). Histological analysis further revealed increased cardiomyocyte cross-sectional area and interstitial fibrosis in csKO hearts (Fig. 3J–L). In addition, expression of the fetal gene markers ANP and βMHC was elevated in PARP16-deficient hearts (Fig. 3M). Collectively, these findings indicate that cardiomyocyte-specific deletion of PARP16 is sufficient to recapitulate the pathological cardiac remodelling phenotype observed in whole-body knockout mice, supporting a cell-autonomous role for PARP16 in maintaining cardiac homeostasis.

**Figure 3.**
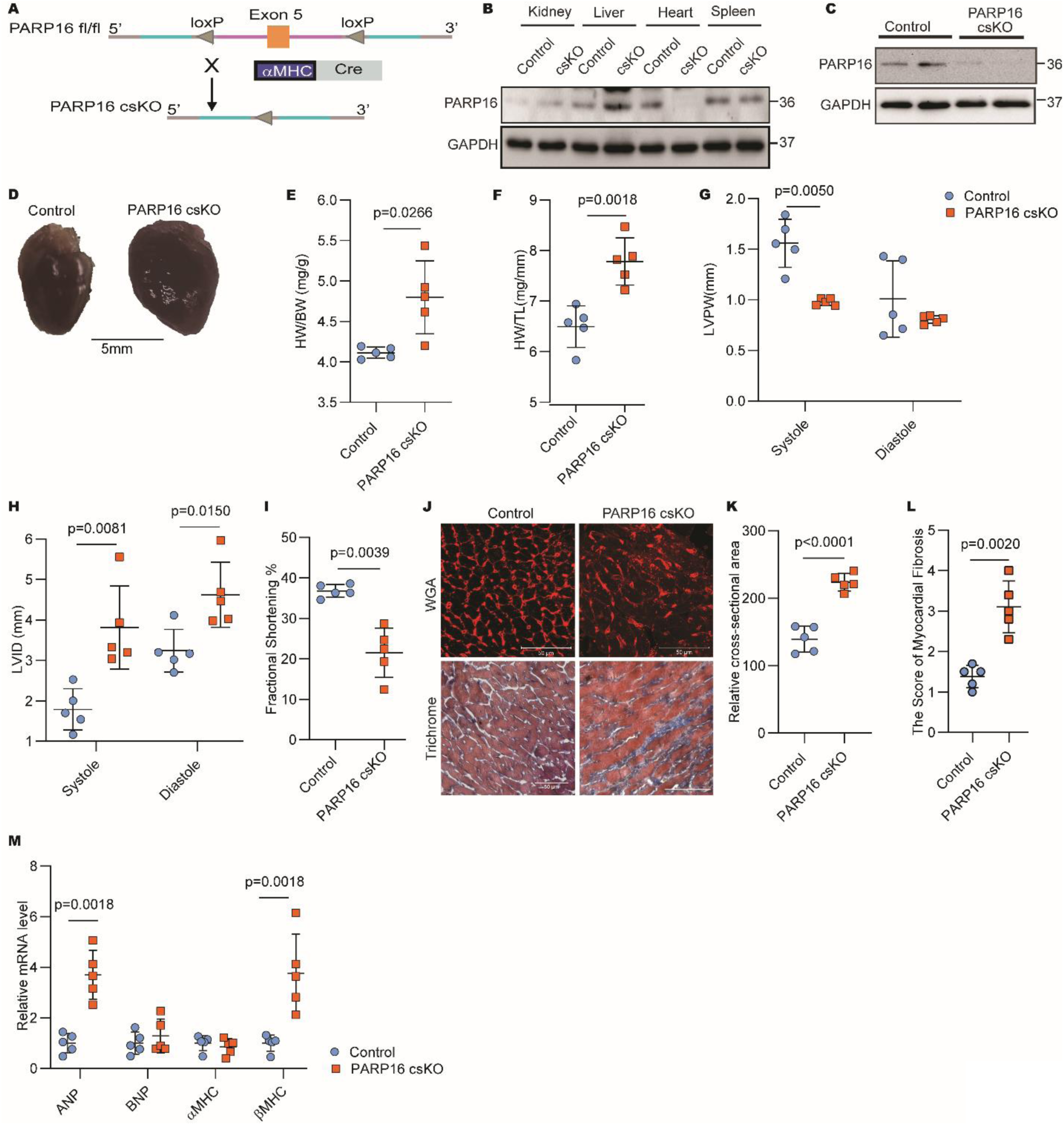
Cardiomyocyte-specific deletion of PARP16 is sufficient to drive pathological cardiac remodelling. (A) Schematic illustrating the generation of cardiomyocyte-specific PARP16 knockout (csKO) mice by crossing PARP16 floxed mice with αMHC-Cre mice. (B) Representative immunoblot analysis of PARP16 protein expression in multiple tissues from3-month-old control and PARP16 csKO mice. Molecular weight markers are indicated. (C) Representative immunoblot of PARP16 protein expression in heart tissue from 3-month-old control and PARP16 csKO mice. (D) Representative gross morphology of hearts isolated from 3-month-old control and PARP16 csKO mice. (E) Heart weight-to-body weight (HW/BW) ratio in 3-month-old control and PARP16 csKO mice. (F) Heart weight-to-tibia length (HW/TL) ratio in 3-month-old control and PARP16 csKO mice. (G) Left ventricular posterior wall thickness (LVPW) at systole and diastole measured by echocardiography in 3-month-old control and PARP16 csKO mice. (H) Left ventricular internal diameter (LVID) at systole and diastole measured by echocardiography in 3-month-old control and PARP16 csKO mice. (I) Fractional shortening (FS) determined by echocardiography in 3-month-old control and PARP16 csKO mice. (J) Representative wheat germ agglutinin (WGA)-stained heart sections for assessment of cardiomyocyte cross-sectional area and Masson’s trichrome-stained sections for assessment of cardiac fibrosis in hearts from 3-month-old control and PARP16 csKO mice. Scale bar, 50 μm. (K) Quantification of cardiomyocyte relative cross-sectional area from WGA-stained heart sections. (L) Quantification of fibrotic score from Masson’s trichrome-stained heart sections. (M) Quantitative RT-PCR analysis of ANP, BNP, αMHC, and βMHC mRNA expression in hearts from 3-month-old control and PARP16 csKO mice. Data are presented as mean ± SD. Statistical significance was determined using unpaired two-tailed t-tests with Welch’s correction. n = 5 mice per group for panels E–M unless otherwise indicated. For panel C, n = 2 mice per group. Exact p values are provided in the source data. Source data are provided as a Source Data file.

### Cardiomyocyte-specific deletion of PARP16 in adult mice induces pathological cardiac remodelling

To determine whether the cardiac abnormalities observed in constitutive cardiomyocyte-specific PARP16 knockout mice were influenced by developmental effects, we generated tamoxifen-inducible cardiomyocyte-specific PARP16 knockout (icsKO) mice (Fig. 4A–D). PARP16 deletion was induced at 8 weeks of age, and cardiac phenotypes were assessed after efficient recombination. As in the constitutive cardiomyocyte-specific knockout model, PARP16 icsKO mice exhibited increased heart weight-to-body weight (HW/BW) and heart weight-to-tibia length (HW/TL) ratios compared with control littermates (Fig. 4E, F). Echocardiographic analysis revealed reduced left ventricular posterior wall thickness (LVPW), increased left ventricular internal diameter during systole (LVIDs), and decreased fractional shortening (Fig. 4G–I), supporting ventricular dilatation and impaired cardiac function. Histological analysis showed increased cardiomyocyte cross-sectional area and interstitial fibrosis in PARP16 icsKO hearts (Fig. 4J–L). In addition, expression of the fetal gene markers ANP and βMHC was significantly increased following inducible PARP16 deletion (Fig. 4N). Together, these findings indicate that adult-onset loss of PARP16 in cardiomyocytes is sufficient to drive pathological cardiac remodelling. The close phenotypic similarity between inducible and constitutive cardiomyocyte-specific knockout models suggests that this phenotype is not attributable to developmental adaptation and instead reflects a continued requirement for PARP16 in maintaining cardiac homeostasis.

**Figure 4.**
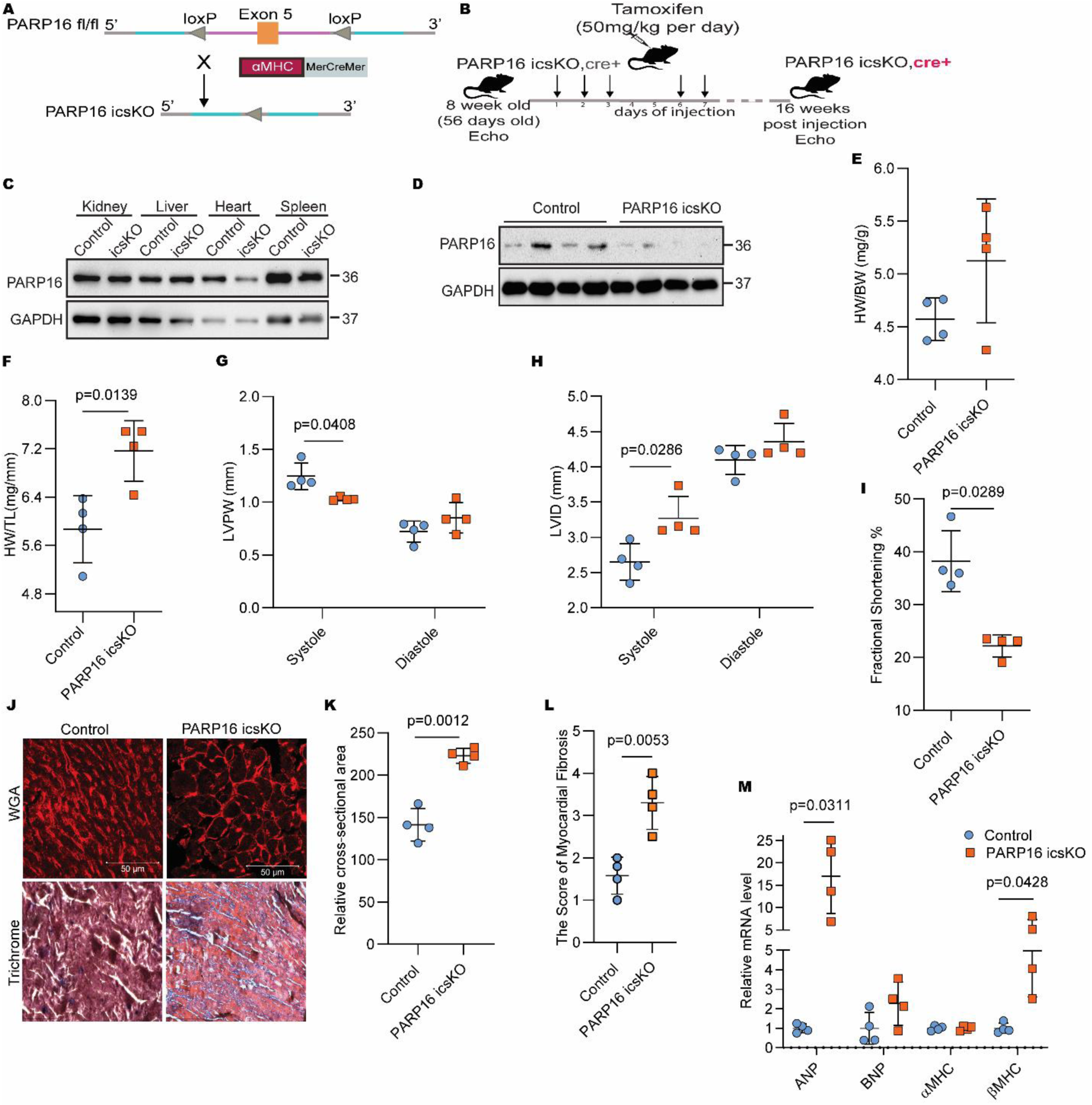
Pathological cardiac remodelling following inducible cardiomyocyte-specific deletion of PARP16. (A) Schematic illustrating the generation of tamoxifen-inducible cardiomyocyte-specific PARP16 knockout (icsKO) mice. (B) Experimental timeline showing tamoxifen administration and phenotypic assessment of control and icsKO mice. (C) Representative immunoblot analysis of PARP16 protein expression in multiple tissues from 6-month-old control and icsKO mice. Molecular weight markers are indicated. (D) Representative immunoblot of PARP16 protein expression in heart tissue from 6-month-old control and icsKO mice following tamoxifen-induced recombination. (E) Heart weight-to-body weight (HW/BW) ratio in 6-month-old control and icsKO mice. (F) Heart weight-to-tibia length (HW/TL) ratio in 6-month-old control and icsKO mice. (G) Left ventricular posterior wall thickness (LVPW) at systole and diastole measured by echocardiography in 6-month-old control and icsKO mice. (H) Left ventricular internal diameter (LVID) at systole and diastole measured by echocardiography in 6-month-old control and icsKO mice. (I) Fractional shortening (FS) determined by echocardiography in 6-month-old control and icsKO mice. (J) Representative wheat germ agglutinin (WGA)-stained heart sections for assessment of cardiomyocyte cross-sectional area and Masson’s trichrome-stained sections for assessment of cardiac fibrosis in hearts from 6-month-old control and icsKO mice. Scale bar, 20 μm. (K) Quantification of cardiomyocyte relative cross-sectional area from WGA-stained heart sections. (L) Quantification of cardiac fibrosis score from Masson’s trichrome-stained heart sections. Data are presented as mean ± SD. Statistical significance was determined using unpaired two-tailed t-tests with Welch’s correction for panels E–G, K, and L, and Mann–Whitney tests for panels H and I, as indicated in the figure. n = 4 mice per group for all quantitative analyses unless otherwise indicated. Exact P values are provided in the source data. Source data are provided as a Source Data file.

### Cardiomyocyte-specific PARP16 overexpression attenuates isoproterenol-induced cardiac remodelling

To determine whether increased PARP16 expression influences cardiac structure and function, we generated cardiomyocyte-specific PARP16 transgenic (csTG) mice (Fig. 5A–C). Under basal conditions, csTG mice displayed heart weight-to-body weight (HW/BW) and heart weight-to-tibia length (HW/TL) ratios comparable to those of control littermates (Fig. 5D, E). Echocardiographic parameters, including left ventricular posterior wall thickness (LVPW), left ventricular internal diameter (LVID), and fractional shortening, were likewise similar between groups (Fig. 5F–H), indicating that increased PARP16 expression does not adversely affect basal cardiac function. To assess the response to pathological stress, control and csTG mice were treated with isoproterenol (ISO). As expected, ISO induced pathological cardiac remodelling in control mice, with marked alterations in cardiac structure and function (Fig. 5I–L). By contrast, csTG mice were largely protected from these ISO-induced changes. Histological analysis further showed reduced cardiomyocyte hypertrophy and fibrosis in ISO-treated csTG hearts relative to ISO-treated control hearts (Fig. 5M–O). These findings indicate that cardiomyocyte-specific PARP16 overexpression does not perturb basal cardiac physiology but attenuates isoproterenol-induced pathological cardiac remodelling.

**Figure 5.**
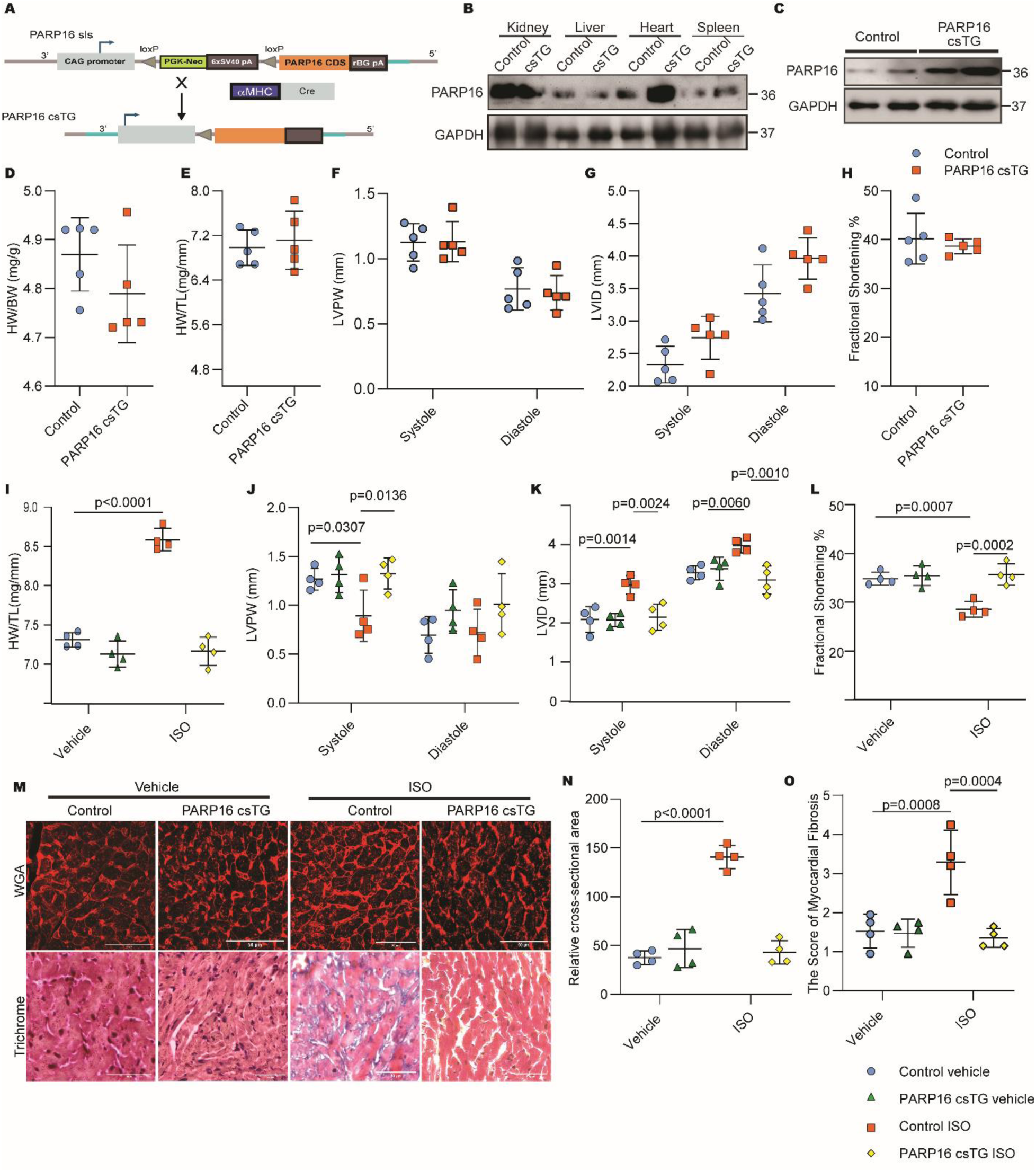
Cardiomyocyte-specific PARP16 overexpression protects against isoproterenol-induced cardiac remodelling. (A) Schematic illustrating the generation of cardiomyocyte-specific PARP16 transgenic (csTG) mice. (B) Representative immunoblot analysis of PARP16 protein expression in multiple tissues from control and csTG mice. Molecular weight markers are indicated. (C) Representative immunoblot of PARP16 protein expression in heart tissue from 3-month-old control and csTG mice. (D) Heart weight-to-body weight (HW/BW) ratio in 3-month-old control and csTG mice. (E) Heart weight-to-tibia length (HW/TL) ratio in 3-month-old control and csTG mice. (F) Left ventricular posterior wall thickness (LVPW) at systole and diastole measured by echocardiography in 3-month-old control and csTG mice. (G) Left ventricular internal diameter (LVID) at systole and diastole measured by echocardiography in 3-month-old control and csTG mice. (H) Fractional shortening (FS) determined by echocardiography in 3-month-old control and csTG mice. (I) Heart weight-to-tibia length (HW/TL) ratio in 3-month-old control and csTG mice following vehicle or isoproterenol (ISO) treatment. (J) Left ventricular posterior wall thickness (LVPW) at systole and diastole measured by echocardiography in 3-month-old control and csTG mice following vehicle or ISO treatment. (K) Left ventricular internal diameter (LVID) at systole and diastole measured by echocardiography in 3-month-old control and csTG mice following vehicle or ISO treatment. (L) Fractional shortening (FS) determined by echocardiography in 3-month-old control and csTG mice following vehicle or ISO treatment. (M) Representative wheat germ agglutinin (WGA)-stained heart sections for assessment of cardiomyocyte cross-sectional area and Masson’s trichrome-stained sections for assessment of cardiac fibrosis in hearts from control and csTG mice following vehicle or ISO treatment. Scale bar, 50 μm. (N) Quantification of cardiomyocyte relative cross-sectional area from WGA-stained heart sections. (O) Quantification of cardiac fibrosis score from Masson’s trichrome-stained heart sections. Data are presented as mean ± SD. Statistical significance was determined using unpaired two-tailed t-tests with Welch’s correction for panels D–H and two-way ANOVA followed by Sidak’s multiple-comparisons test for panels I–L, N, and O, as indicated in the figure. n = 5 mice per group for panels D–H; n = 4 mice per group for panels I–L, N, and O; n = 2 mice per group for panel C. Exact p-values are provided in the source data. Source data are provided as a Source Data file.

### Transcriptomic profiling identifies several pathways involved in the cardiac remodelling of PARP16 deficient mice

To investigate the molecular mechanisms underlying the remodelling phenotype associated with PARP16 deficiency, we performed RNA sequencing on hearts from control and PARP16 knockout mice (Fig. 6A). Differential expression analysis revealed extensive transcriptional changes following PARP16 deletion, including enrichment of gene signatures associated with pathological cardiac remodelling and dilated cardiomyopathy (Fig. 6B, C). A heatmap of differentially expressed genes further demonstrated broad transcriptomic reprogramming in PARP16-deficient hearts (Fig. S1A). To identify transcriptional regulators that may contribute to these changes, we performed Integrated System for Motif Activity Response Analysis (ISMARA). This analysis predicted altered activity of several transcription factor families, including NFAT, GATA4, and MEF2 (Fig. 6E, F and Fig. S1C, D). These findings suggest the possible involvement of NFAT-GATA4-MEF2 transcriptional factors in the pathological cardiac remodelling observed in the PARP16-KO mice. Given the established role of PARP16 in unfolded protein response signalling, we next examined canonical UPR markers. Immunoblotting and gene expression analyses did not reveal marked alterations in the major UPR pathways in PARP16-deficient hearts (Fig. S1E-G). These findings suggest that the transcriptomic changes observed following PARP16 deficiency are not accompanied by overt activation of canonical UPR signalling under the conditions examined. Therefore, we next investigated whether NFAT signalling, a pathway strongly linked to pathological cardiac remodelling, is functionally altered following PARP16 deficiency.

**Figure 6.**
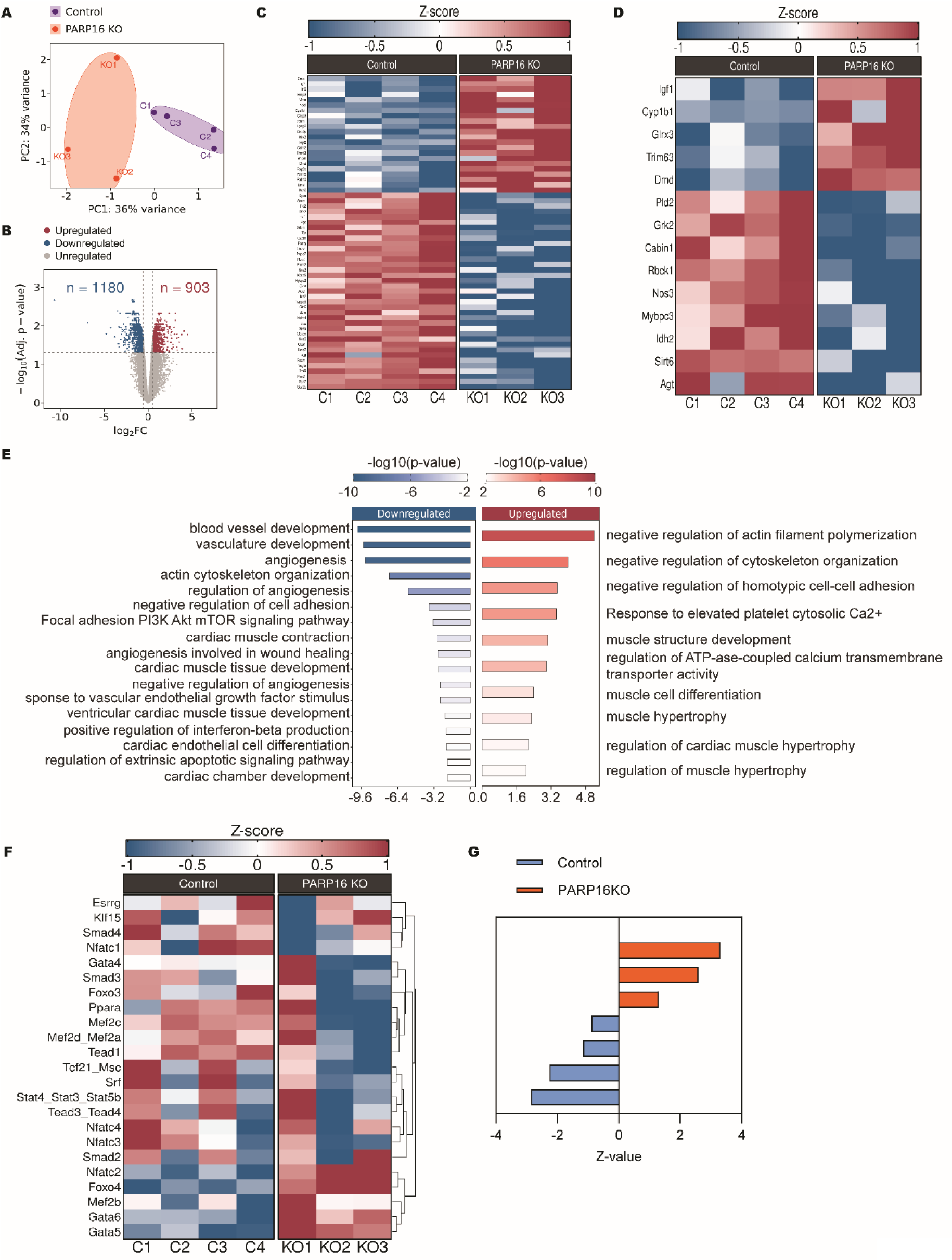
RNA-sequencing and motif activity analyses reveal NFAT pathway activation in PARP16-deficient hearts. (A) Principal component analysis (PCA) of RNA sequencing data from hearts of 5-month-old control (n = 4) and PARP16 knockout (KO) mice (n = 3). (B) Volcano plot showing differentially expressed genes identified by RNA sequencing in hearts from 5-month-old control and PARP16 KO mice. (C) Heatmap of differentially expressed genes associated with dilated cardiomyopathy-related genes in5-month-old 5-month-old control and PARP16 KO hearts. (D) Heatmap of differentially expressed genes associated with cardiac remodelling and hypertrophy-related genes in 5-month-old control and PARP16 KO hearts. (E) Gene Ontology Biological Process (GO-BP) enrichment analysis of selected upregulated and downregulated biological processes identified from RNA sequencing data. (F) Heatmap of predicted transcription factor activities generated using Integrated System for Motif Activity Response Analysis (ISMARA) from 5-month-old control and PARP16 KO heart transcriptomes. (G) Predicted NFAT1 transcription factor activity scores derived from ISMARA analysis for individual control and PARP16 KO heart samples. Data shown in panels A–G were generated from RNA sequencing analysis of heart tissue isolated from 5-month-old control and PARP16 KO mice. Statistical methods used for differential expression and enrichment analyses are described in the Methods section. Source data are provided as a Source Data file.

### PARP16 deficiency promotes pathological cardiac remodelling through NFAT signalling

To validate the transcriptomic prediction of enhanced NFAT signalling, we first assessed the expression of established NFAT target genes in PARP16 knockout hearts. Quantitative RT-PCR revealed significantly increased expression of Pik3r1, Agt, Itpr2, Adssl1, Slim1, and Atp2a2 compared with control hearts (Fig. 7A). Immunoblot analysis further demonstrated elevated NFAT1 protein levels in PARP16-deficient hearts (Fig. 7B and Fig. S2A). As the transcriptional activity of NFAT is governed by its subcellular localization, we next evaluated the distribution of NFAT1 in control and PARP16 knockout hearts. Nuclear-cytoplasmic fractionation revealed increased nuclear accumulation of NFAT1 accompanied by a corresponding reduction in the cytoplasmic fraction (Fig. 7C, D). ChIP-qPCR further demonstrated increased NFAT1 occupancy at the promoters of Rcan1, βMHC, and Acta1 in PARP16-deficient hearts (Fig. 7E), supporting enhanced NFAT-dependent transcription.

**Figure 7.**
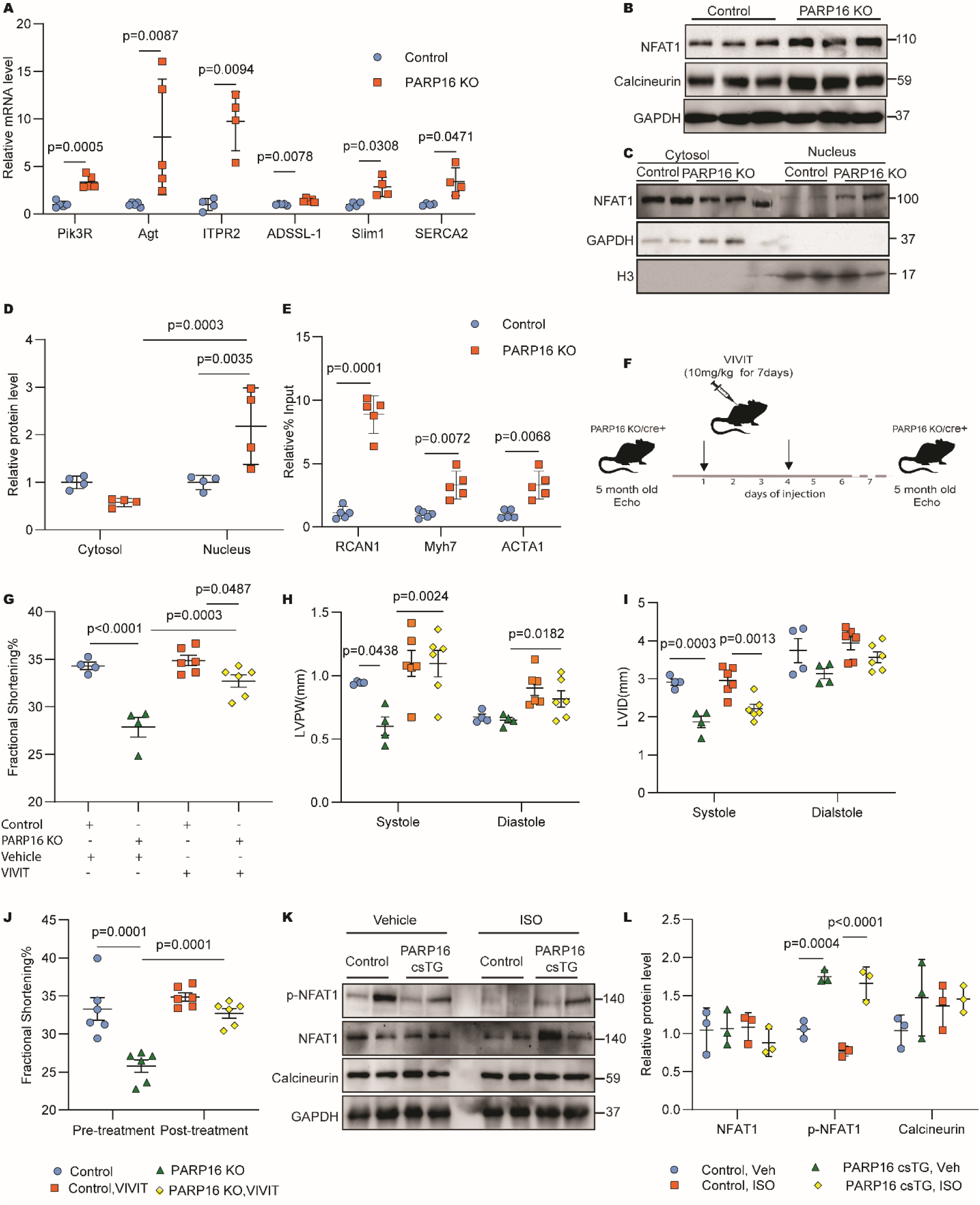
Activation of NFAT signalling contributes to pathological cardiac remodelling following PARP16 deficiency. (A) Quantitative RT-PCR analysis of NFAT1 target gene expression in hearts from 5-month-old control and PARP16 knockout (KO) mice (n = 4–5 mice per group). (B) Representative immunoblot analysis of NFAT1 and calcineurin protein expression in heart tissue from 5-month-old control and PARP16 KO mice (n = 3 mice per group). (C) Representative immunoblot analysis of NFAT1 protein expression in cytosolic and nuclear fractions isolated from hearts of 5-month-old control and PARP16 KO mice (n = 2 mice per group). (D) Quantification of NFAT1 protein abundance in cytosolic and nuclear fractions. GAPDH and Histone H3 were used as cytosolic and nuclear loading controls, respectively. (E) Chromatin immunoprecipitation (ChIP)-qPCR analysis of NFAT1 occupancy at selected target gene promoters in hearts from 5-month-old control and PARP16 KO mice (n = 5 mice per group). (F) Schematic illustrating the VIVIT treatment regimen used for pharmacological inhibition of NFAT signalling *in vivo*. (G) Fractional shortening (FS) measured by echocardiography in 5-month-old control and PARP16 KO mice treated with vehicle or VIVIT. (H) Left ventricular posterior wall thickness (LVPW) at systole and diastole measured by echocardiography in 5-month-old control and PARP16 KO mice treated with vehicle or VIVIT. (I) Left ventricular internal diameter (LVID) at systole and diastole measured by echocardiography in 5-month-old control and PARP16 KO mice treated with vehicle or VIVIT. (J) Fractional shortening measured before and after VIVIT treatment in PARP16 KO mice. (K) Representative immunoblot analysis of NFAT1 signalling components in hearts from 3-month-old isoproterenol (ISO)-treated control and cardiomyocyte-specific PARP16 transgenic (csTG) mice (n = 2 mice per group). (L) Quantification of phosphorylated NFAT1, total NFAT1, and calcineurin protein abundance in hearts from 3-month-old ISO-treated control and csTG mice. Protein levels were normalised to GAPDH. Data are presented as mean ± SD. Statistical analyses were performed as indicated in the figure. Exact P values are provided in the source data. Source data are provided as a Source Data file.

To investigate whether enhanced NFAT signalling contributes to the remodelling phenotype, PARP16 knockout mice were treated with the selective NFAT inhibitor VIVIT. VIVIT treatment significantly improved fractional shortening and attenuated abnormalities in ventricular wall thickness and chamber dimensions compared with vehicle-treated PARP16 knockout mice (Fig. 7G-J). Similar effects were observed in PARP16-deficient neonatal rat cardiomyocytes, where VIVIT reduced perinuclear ANP accumulation, improved sarcomeric organization (Fig. S2B, C), and attenuated the increase in NFAT-dependent luciferase activity induced by PARP16 depletion (Fig. S2D). These findings demonstrate that enhanced NFAT signalling is required for the pathological remodelling associated with PARP16 deficiency. To support the findings of these loss-of-function studies, we next assessed NFAT signalling in isoproterenol-treated cardiomyocyte-specific PARP16 transgenic hearts. PARP16 overexpression preserved phosphorylated NFAT levels following isoproterenol treatment without altering total NFAT1 protein abundance (Fig. 7K, L), further supporting suppression of NFAT signalling by PARP16 under pathological stress. Together, these findings establish enhanced NFAT signalling as a key mediator of the pathological remodelling induced by PARP16 deficiency.

### PARP16 suppresses NFAT1 transcriptional activity via ADP-ribosylation

Having established enhanced NFAT signaling as a key mediator of the remodeling phenotype associated with PARP16 deficiency, we next investigated the molecular mechanism by which PARP16 regulates NFAT1. NFAT luciferase assays showed that overexpression of wild-type PARP16 significantly reduced NFAT transcriptional activity under both basal conditions and following PARP16 depletion (Fig. 8A). In contrast, the catalytically inactive PARP16 mutant (Y254A) failed to suppress NFAT activity in PARP16-deficient cells, indicating that the enzymatic activity of PARP16 is required for inhibition of NFAT1 signalling. To assess whether PARP16 influences NFAT-dependent transcription, chromatin immunoprecipitation was performed at the Rcan1 and βMHC promoters. Wild-type PARP16 significantly reduced NFAT1 occupancy at both promoters, whereas the PARP16 catalytic mutant increased the NFAT1 occupancy (Fig. 8B, C), indicating that PARP16 suppresses NFAT-dependent transcription in an enzymatic activity-dependent manner.

**Figure 8.**
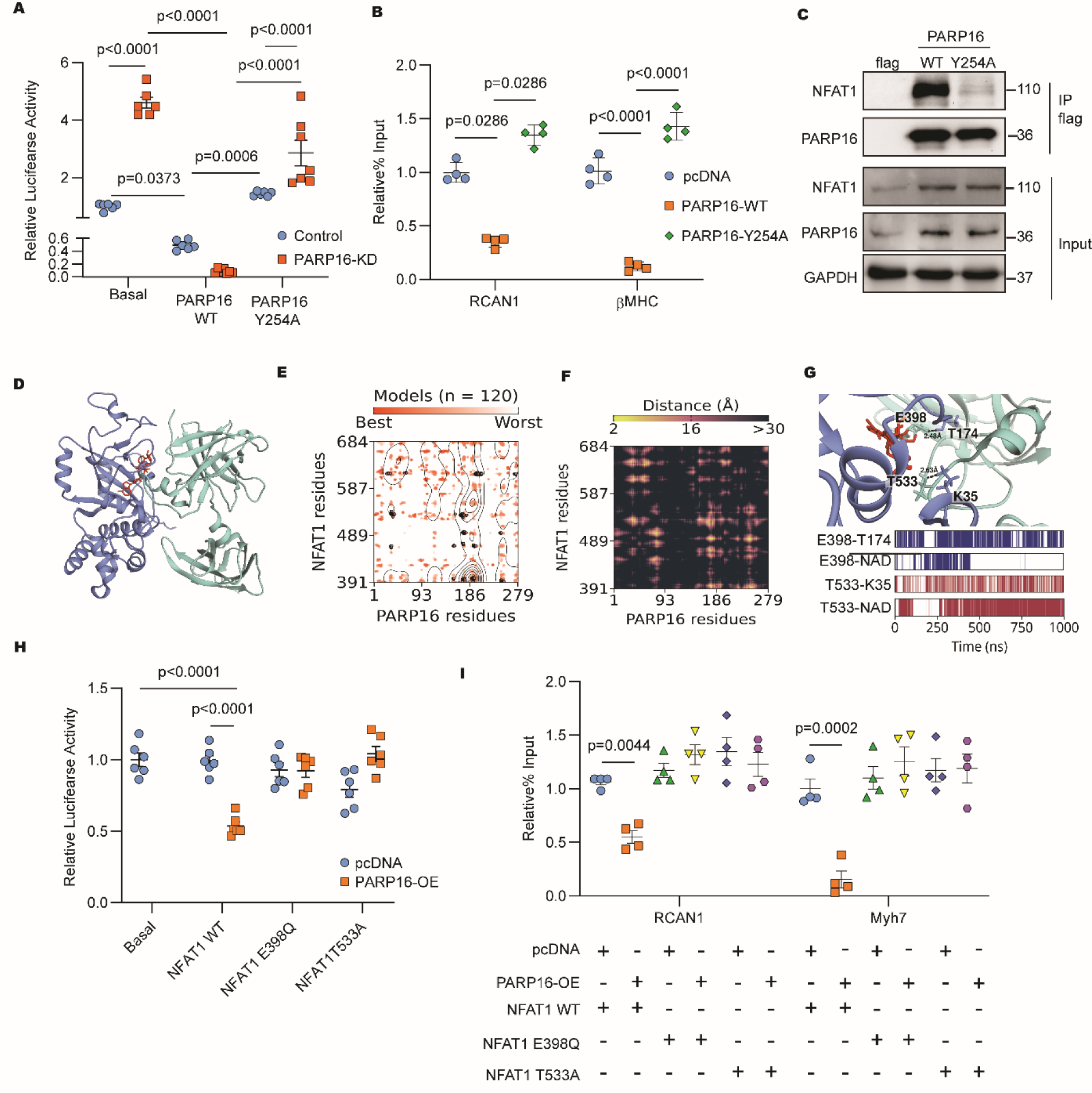
PARP16 interacts with NFAT1 and candidate NFAT1 residues contribute to PARP16-mediated regulation of NFAT1 activity. (A) NFAT1 luciferase reporter activity in neonatal rat cardiomyocytes (NRCMs) expressing PARP16 wild-type (WT) or catalytic mutant (Y254A) under control or PARP16 knockdown conditions. (B) Chromatin immunoprecipitation (ChIP)-qPCR analysis of NFAT1 occupancy at selected target gene promoters in cells expressing PARP16 WT or PARP16 Y254A. (C) Co-immunoprecipitation analysis of PARP16 and endogenous NFAT1 in NRCMs following overexpression of PARP16-FLAG or PARP16 Y254A-FLAG. Immunoprecipitation was performed using anti-FLAG magnetic beads and immunoblots were probed for NFAT1. (D) Top-ranked docking model of the PARP16–NAD⁺–NFAT1 complex generated using ClusPro. PARP16 is shown in dark blue, NFAT1 in cyan, and NAD⁺ in red. (E) Distribution of PARP16–NFAT1 interfacial residue pairs identified within 8 Å across 120 ClusPro-generated docking models. Residue pairs present in the top-ranked model are highlighted. (F) Location of candidate NFAT1 residues E398 and T533 identified by mass spectrometry and selected for subsequent structural and functional analyses. (G) Molecular dynamics analysis of the PARP16–NAD⁺–NFAT1 complex. Top, average structure derived from a 1000-ns simulation showing the relative positioning of NFAT1 residues E398 and T533 with respect to PARP16 and NAD⁺. Bottom, interaction profiles observed throughout the simulation trajectory, including hydrogen-bonding and van der Waals interactions involving E398, T533, PARP16 residues, and NAD⁺. (H) NFAT1 luciferase reporter activity in cells expressing NFAT1 WT, E398Q, T533A, or the indicated mutant constructs under basal or PARP16-overexpressing conditions. (I) ChIP-qPCR analysis of NFAT1 occupancy at selected target gene promoters in cells expressing NFAT1 WT, E398Q, or T533A under basal or PARP16-overexpressing conditions. Data are presented as mean ± SD. Statistical analyses were performed as indicated in the figure. Exact P values are provided in the source data. Source data are provided as a Source Data file.

We next evaluated whether PARP16 physically interacts with NFAT1. Co-immunoprecipitation experiments demonstrated interaction between PARP16 and NFAT1 (Fig. 8D). To gain structural insights into this interaction, refined structural models of PARP16 and NFAT1 were generated. The crystal structure of human PARP16 (PDB ID: 6HXS) served as the template, with unresolved regions modelled using AlphaFold2 [34]. For NFAT1, the crystal structure encompassing the Rel homology domain (PDB ID: 1A02) was similarly completed using AlphaFold2 [33]. The PARP16-NAD complex was generated using AlphaFold3 [34], followed by protein-protein docking using HDOCK[35], LightDock[36], and LZerD[37]. Independent docking approaches converged on a similar interaction interface, positioning the catalytic domain of PARP16 adjacent to the NFAT1 Rel homology domain (Fig. S3A-E).

To identify residues potentially modified by PARP16, NFAT1 was analysed by LC-MS/MS following co-expression with wild-type or catalytically inactive PARP16. ADP-ribosylated NFAT1 peptides were reproducibly detected only in samples expressing wild-type PARP16 and were absent from catalytic mutant and control samples (Fig. 8E and Fig. S3F). We identified two residues, E398 and T533, were consistently ADP-ribosylated across three biological replicates. To further evaluate the structural relevance of these candidate residues, 1000-ns all-atom molecular dynamics simulations were performed using the PARP16-NAD-NFAT1 complex. PARP16 remained stably associated with NFAT1 throughout the simulation (Fig. 8F). Distance analyses positioned NFAT1 E398 in proximity to PARP16 T174 and NFAT1 T533 near PARP16 K35, with both residues located adjacent to the bound NAD molecule (Fig. 8G). Notably, E398 and T533 reside within the NFAT1 Rel homology domain responsible for DNA binding and transcriptional regulation.

To assess the functional significance of these candidate residues, NFAT1 E398Q and T533A mutants were generated and evaluated using NFAT luciferase assays. Unlike wild-type NFAT1, neither mutant underwent transcriptional suppression following PARP16 overexpression (Fig. 8H). Similarly, PARP16 overexpression reduced NFAT1 occupancy at the Rcan1 and βMHC promoters in cells expressing wild-type NFAT1, whereas this effect was markedly attenuated in cells expressing either mutant (Fig. 8I). These findings demonstrate the functional importance of E398 and T533 in PARP16-dependent regulation of NFAT1. Collectively, these data demonstrate that PARP16 suppresses NFAT1 transcriptional activity through its enzymatic function and identify E398 and T533 as candidate residues that contribute to this regulatory mechanism.

## Discussion

Pathological cardiac remodelling is driven by coordinated changes in intracellular signalling, transcriptional regulation, and post-translational modifications that ultimately culminate in ventricular dysfunction and heart failure [13–16]. Although PARP1 and PARP2 have established roles in cardiovascular disease [14, 19–22], considerably less is known about the physiological functions of mono-ADP-ribosyltransferases in the heart [16–18]. The present study identifies PARP16 as a previously unrecognised regulator of cardiac homeostasis. Using constitutive, cardiomyocyte-specific, and inducible loss-of-function models together with cardiomyocyte-specific overexpression, we demonstrate that reduced PARP16 expression promotes adverse cardiac remodelling, whereas increased PARP16 expression attenuates stress-induced remodelling. Mechanistically, our data support a model in which PARP16 reduces NFAT1-dependent transcription activity through a catalytic activity-dependent mechanism. The translational relevance of these findings is supported by reduced PARP16 expression in failing human hearts and in an independent transcriptomic dataset from patients with dilated cardiomyopathy. Similar remodelling phenotypes observed in constitutive, cardiomyocyte-specific, and inducible knockout mice indicate that the cardiac abnormalities are unlikely to arise from developmental adaptation. Together, these observations suggest that loss of PARP16 may contribute to disease progression in the failing heart.

PARP16 has primarily been studied as a regulator of PERK- and IRE1α-dependent unfolded protein response signalling [23–25]. This suggested that altered UPR signalling is a plausible mechanism underlying the phenotype observed in PARP16-deficient hearts. However, assessment of canonical UPR markers did not reveal overt alterations under the conditions examined. Instead, unbiased transcriptomic analyses identified NFAT signalling as a candidate pathway associated with PARP16 deficiency. Subsequent molecular, biochemical, and pharmacological studies consistently supported enhanced NFAT signalling, including increased expression of NFAT-responsive genes, enhanced nuclear accumulation of NFAT1, increased promoter occupancy, and attenuation of the remodelling phenotype following NFAT inhibition. These observations identify NFAT signalling as a principal mediator of the pathological remodelling associated with PARP16 deficiency, while not excluding additional PARP16-regulated pathways under different pathological settings.

A recent study implicated PARP16 in pathological cardiac remodelling by activating the IRE1α-sXBP1-GATA4 signalling axis [26]. Our findings instead identify NFAT1 as the principal downstream effector involved in cardiac remodelling. These findings could be due to the differences in disease models and experimental approaches. Whereas the previous study primarily employed AAV9-mediated knockdown and pressure-overload models, the present study combined multiple genetic mouse models with transcriptomic profiling, pharmacological intervention, and mechanistic analyses. These differences may account for the distinct downstream pathways identified and support the view that PARP16 regulates stress-responsive signalling in a context-dependent manner.

NFAT transcription factors are central regulators of pathological cardiac remodelling and heart failure [27, 38, 39]. Previous studies demonstrated that PARP1-mediated ADP-ribosylation enhances NFAT transcriptional activity and DNA binding [40, 41], whereas our data identify PARP16 as a negative regulator of NFAT1-dependent transcription. These observations are not mutually exclusive because PARP1 and PARP16 differ substantially in catalytic activity, intracellular localisation, and substrate specificity [17, 18, 23]. Our findings therefore suggest that the functional consequences of ADP-ribosylation on NFAT signalling depend on the modifying PARP family member and the biological context. Mechanistically, we identified E398 and T533 as candidate residues associated with PARP16-dependent regulation of NFAT1. Although the intrinsic lability of ADP-ribosylation precludes unequivocal assignment of modification sites by conventional LC-MS/MS, these candidate residues were reproducibly detected only in the presence of catalytically active PARP16 and were further supported by structural modelling and functional mutagenesis. Their localization within the NFAT1 Rel homology domain further supports a role in regulating NFAT transcriptional function.

A few limitations should be acknowledged in this study. Although our data establish NFAT1 as a major downstream mediator of PARP16 in pathological cardiac remodelling, they do not exclude the contribution of additional PARP16-regulated signalling pathways under different pathological conditions. We have subjected the PARP-16 transgenic mice to cardiac stress induced by Isoproterenol; however, the effect of PARP16 activation needs to be tested in other models of cardiac stress such as pressure overload. Despite these limitations, the present work identifies a previously unrecognized PARP16-NFAT1 signalling axis that protects against pathological cardiac remodelling and provides a mechanistic framework for understanding how mono-ADP-ribosylation regulates stress-responsive transcription in the heart.

## RESOURCE AVAILABILITY

### Lead Contact

Further information and requests for resources and reagents should be directed to the Corresponding Author, N. Ravi Sundaresan.

### Materials availability

All unique/stable reagents generated in this study are available with the Lead Contact (N. Ravi Sundaresan with a completed Materials Transfer Agreement.

### Data availability

RNA sequencing data are available and deposited in the GEO database with accession number **PRJNA1425995**. Mass spectroscopy data are available and deposited in the PRIDE database with accession number **PXD074616**. The datasets generated and analysed during the current study are available from the lead contact upon reasonable request.

## SUPPLEMENTAL MATERIAL

For information regarding reagent specifications, refer to the Major Resources Table in Supplementary File 1.

For Source data related to figures used in this study, please refer to Supplementary file 2.

## ACKNOWLEDGEMENTS

We acknowledge the Central Animal Facility, Central Confocal facility (Division of Biological Sciences and Department of Microbiology and Cell Biology), and the microtome facility (Division of Biological Sciences) at IISc, Bengaluru, for their services and technical help. S.Z. acknowledges IISc for providing a fellowship to support this study.

## AUTHOR CONTRIBUTIONS

N.R.S. conceptualisation, project administration, and supervision, manuscript review, and editing. S.Z. conceptualised, designed and analysed the experiments and wrote the first draft of the manuscript. S.R. experimental design, scientific discussion, methodology, animal work. A.B.P. methodology for mass spec data, Scientific discussion. A.C. writing, Simulation of mass spectrometry data and RNA-seq data analysis. P.J. data quantification. R.J. human sample collection. A.S. RNA-seq data Analysis. AJ.S. manuscript writing and editing. P.A.R. data quantification. M.H. conceptualisation, writing, review, and editing. K.S. conceptualisation. U.N. conceptualisation. P.S.M.R. providing human patient samples.

## Materials and methods

### Human heart samples

All research involving human samples was conducted in accordance with the Institutional Human Ethics Committee approval (IHEC No: 06/22.06.2023) and complied with institutional and national ethical guidelines. Inclusion and exclusion criteria for sample collection were as previously described [42]. Written informed consent was obtained from all living participants or from their legally authorised representatives for the donation of tissues. Left ventricular myocardial tissue was obtained from patients undergoing cardiac surgery at Sri Jayadeva Institute of Cardiovascular Sciences and Research, Bengaluru, India. Clinical evaluation, including cardiac function assessment, was performed according to established AHA/ACC, European, and Indian guidelines. Recruitment was based on MRI and echocardiographic reports as described in[42]. Patients with advanced left ventricular dysfunction were included. Approximately 100 mg of tissue from the left ventricular free wall was collected intraoperatively using an endomyocardial punch biopsy technique. Non-failing control left ventricular tissue was obtained from deceased male donors without documented cardiovascular disease. Control samples were derived from the left ventricular free wall. Patients from whom adequate biological material or essential clinical data were unavailable, or who did not consent to research use, were excluded. All specimens were snap-frozen in liquid nitrogen immediately after procurement and stored at −80°C until analysis. Clinical characteristics of the analysed samples are summarised in Supplementary Table 2.

### Generation of PARP16 knock-out and transgenic mice models

All animal experiments were conducted in accordance with the ethical guidelines established by the Committee for the Purpose of Control and Supervision of Experiments on Animals (CPCSEA), Government of India, and were approved by the Institutional Animal Ethics Committee (CAF/Ethics/885/2022). To minimise variability associated with sex-specific hormonal influences, only male mice were used in all experiments. Mice were housed in individually ventilated cages under a 12-hour light/dark cycle with ad libitum access to standard chow and water. PARP16 Flox⁺/⁻ mice (Cyagen Biosciences, China) were intercrossed to generate homozygous PARP16 Flox⁺/⁺ mice. Genotyping was performed by PCR using specific primers (Forward: 5′-AAGGTCAGATGCTACAGGTAGGA-3′; Reverse: 5′-CAGCAAACACTCCTGACCGCTAG-3′). PCR conditions consisted of initial denaturation at 95°C for 3 minutes, followed by 35 cycles of 94°C for 30 seconds, 60°C for 35 seconds, and 72°C for 35 seconds, with a final extension at 72°C for 5 minutes. PCR products were resolved by agarose gel electrophoresis (157 bp wild-type allele; 224 bp floxed allele).

To generate cardiac-specific knockout (csKO) mice, homozygous PARP16 Flox⁺/⁺ mice were crossed with Myh6-Cre mice (αMHC-Cre). Offspring heterozygous for both the floxed allele and Cre transgene were identified by PCR using Cre-specific primers (Forward: 5′-GAACGCACTGATTTCGACCA-3′; Reverse: 5′-GCTAACCAGCGTTTTCGTTC-3′; 204 bp product). These mice were further bred to obtain the desired experimental genotypes. Littermate controls lacking Cre were used for all comparisons. To generate tamoxifen-inducible cardiac-specific knockout (icsKO) mice, PARP16 Flox⁺/⁺ mice were crossed with αMHC-MerCreMer transgenic mice (Myh6-MerCreMer). Progeny carrying both floxed PARP16 alleles and the MerCreMer transgene were identified by PCR as described above. For induction of recombination, male mice at 8 weeks of age received tamoxifen (Sigma-Aldrich) dissolved in corn oil (10 mg/mL) administered intraperitoneally at 40 mg/kg body weight once daily for 5 consecutive days. Control littermates received vehicle injections. Mice were allowed to recover for 8 weeks following tamoxifen administration before phenotypic assessment. Successful recombination and PARP16 depletion in cardiac tissue were confirmed at the protein level prior to analysis. To generate cardiac-specific Transgenic (csTG) mice, heterozygous PARP16 LsL⁺/⁻ mice (Cyagen Biosciences) were intercrossed to obtain homozygous PARP16 LsL⁺/⁺ mice. Genotyping was performed using allele-specific primer sets (519 bp transgenic allele; 453 bp wild-type allele). Homozygous LsL⁺/⁺ mice were crossed with αMHC-Cre mice to generate cardiac-specific PARP16 overexpression in Cre-positive offspring. Littermate Cre-negative mice were used as controls.

### Primary cardiomyocyte culture

Primary cardiomyocytes were derived from neonatal rat pups and cultured as described in our previous publication [43]. Briefly, the pups were anaesthetised using isoflurane and then sacrificed. The hearts were harvested into PBS-glucose. The collected heart tissue was minced and digested using collagenase II and 0.2% trypsin. At the end of each digestion cycle, the cell pool was collected in horse serum. Upon complete digestion of heart tissue, the cell pool was pre-plated for 45 minutes to separate the fibroblasts from the cardiomyocytes. After the incubation period, the unattached cells (cardiomyocytes) were harvested from the dishes, centrifuged, and then seeded into 0.2% gelatin-coated well plates.

### Cell culture

HEK293 cell line was maintained in Dulbecco’s Modified Eagle Medium (DMEM) supplemented with 10% foetal bovine serum (FBS) and 1X anti-anti cocktail. Cell cultures were incubated at 37°C in a humidified chamber containing 5% CO₂. PARP16 knockdown was performed by transfecting cells with PARP16-specific siRNA using RNAi MAX (Thermo Fisher Scientific) according to the manufacturer’s instructions. Cells were incubated for 72 hours post-transfection. PARP16 overexpression was performed by transfecting cells with the hPARP16 plasmid using Lipofectamine 3000 (Thermo Fisher Scientific) according to the manufacturer’s instructions. Cells were incubated for 48 hours post-transfection. Western blotting and qPCR were performed to experimentally validate both knockdown and overexpression of PARP16.

### Echocardiography analysis

Cardiac function was assessed non-invasively by a trained operator using the VisualSonics Vevo 1100 imaging system (FUJIFILM VisualSonics) equipped with a high-frequency ultrasound transducer in accordance with ASE guidelines for small animals, as described previously[38, 44]. Briefly, mice were anaesthetised with 1% isoflurane, maintaining physiological stability by monitoring body temperature at 37.0±0.5°C and heart rate between 400 and 500 beats per minute. Image acquisition was performed with two-dimensional (2D) guidance in the Parasternal Long-Axis (PLAX) and Parasternal Short-Axis (PSAX) views. Echocardiographic images were analysed using Vevo LAB software (FUJIFILM VisualSonics).

### Histology analysis

Heart tissue was fixed for 72 hours in 10% neutral buffered formalin. The fixed tissue was processed, embedded in paraffin, and sectioned using the microtome to obtain sections of 5 μm thickness. Cell size was measured using wheat germ agglutinin (WGA) stain by standard techniques. Images were acquired using a Zeiss LSM 710 or 880 confocal microscope, followed by quantification using NIH ImageJ software.

### Confocal microscopy analysis

Samples were prepared for confocal microscopy as described previously [45]. Briefly, cardiomyocytes were seeded onto sterile glass coverslips in a 24-well plate and fixed using a chilled mixture of acetone and methanol (1:1) for 15 minutes at −20°C. Following fixation, cells were mildly permeabilized using 0.1% saponin in 1% bovine serum albumin (BSA) for 1 hour at room temperature. Subsequently, cells were incubated overnight at 4°C with specific primary antibodies diluted in 0.1% saponin in 1% BSA. After washing with 1× phosphate-buffered saline (PBS), cells were incubated for 1 hour at room temperature with appropriate secondary antibodies conjugated to Alexa Fluor 488 and Alexa Fluor 546, prepared in 0.1% saponin in 1% BSA. Nuclei were counterstained with Hoechst 33342 for 10 minutes at room temperature. Cells were then washed with 1× PBS and mounted onto clean glass slides using Fluoromount G mounting medium. Images were captured using a Zeiss LSM 710 or 880 confocal microscope.

### RNA isolation and Real-time PCR analysis

Cultured cells and mouse heart tissue were homogenised, and RNA was isolated using TAKARA RNAiso Plus according to the manufacturer’s protocol. 500 ng of the total isolated RNA was used for cDNA synthesis, performed using the PhiScript™ cDNA Synthesis Kit (dx/dt, Bangalore, KA, India) as per the manufacturer’s protocol, and qPCR was performed using TB Green™ Premix Ex Taq™ II (Tli RNaseH Plus) (Takara Bio Inc., Kusatsu, Japan) on a QuantStudioTM 6 Flex System. Thermal cycling conditions were as follows: initial denaturation at 95°C for 30 seconds, followed by 40 cycles of 95°C for 5 seconds and 60°C for 34 seconds. A melting curve analysis was performed to verify amplification specificity. Relative gene expression was calculated using the ΔCt method, with GAPDH as the internal control.

### Immunoblotting

Frozen mouse tissues and cell cultures were homogenised and lysed using RIPA lysis buffer (1 mM EDTA, 2.5 mM sodium pyrophosphate, 5mM nicotinamide, 150 mM NaCl, 20 mM Tris–HCl, pH 7.5, 1mM PMSF, 1mM sodium orthovanadate, 1mM EGTA, 1% NP-40, 1X protease inhibitor cocktail, and 1% sodium deoxycholate). The lysate was vortexed for 30 minutes at 4°C and then centrifuged for 15 minutes at 12,000 rpm, 4°C. Protein concentrations in lysates were determined using the Bradford assay, following the manufacturer’s instructions. Protein estimation was performed using Bradford reagent. Samples were prepared accordingly and then heated at 95°C for 5 minutes in 2X Laemmli buffer supplemented with 5% β-mercaptoethanol for denaturation. 12% SDS-PAGE separated proteins and subsequently transferred to 0.45 mm PVDF membranes via overnight wet transfer (20V). Membranes were blocked with 5% skimmed milk in TBST. Target proteins were detected using the primary antibodies listed in Supplementary File 1. An enhanced chemiluminescence substrate was applied to the membrane, and chemiluminescent signals were detected using a Bio-Rad ChemiDoc. Protein expression analysis and image processing were carried out using Image Lab (Bio-Rad) software.

### Co-immunoprecipitation assay

NRCMs were transfected with FLAG-tagged PARP16, HA-tagged full-length NFAT1, or empty vector plasmids using Lipofectamine™ 3000 Transfection Reagent (Invitrogen, USA), following the manufacturer’s protocol. After 48 hours, cells were harvested using digitonin-based fractionation as previously described. 1000 µg protein was incubated overnight at 4°C with 1 µg of anti-HA antibody on a rotator at 5 rpm. 20 µL magnetic beads were washed three times with digitonin buffer for 30 seconds at 1000 rpm. The lysate protein was added to the washed beads and incubated on a rotator at 5 rpm, 4°C for 4 hours. Beads were washed three times with ice-cold PBS, and bead-bound proteins were eluted by adding 2× Laemmli sample buffer (Bio-Rad) and boiling at 95°C for 5 minutes. Eluted proteins were subjected to immunoblotting.

### RNA Sequencing and Differential Expression Analysis

Raw FASTQ files were quality-assessed using FastQC (v0.11.9; http://www.bioinformatics.babraham.ac.uk/projects/fastqc/) with default parameters. Preprocessing was performed using Fastp (v0.20.1)[58] with the following parameters: --trim_front1 8 --trim_front2 6 --length_required 50 --correction --trim_poly_g --qualified_quality_phred 30. Post-preprocessing quality was reassessed using FastQC, and results were summarised with MultiQC (v1.25.2)[59]. Processed reads were aligned to the Mus musculus GRCm39 genome using STAR aligner (v2.7.9a)[60] with the parameters: --outSAMtype BAM SortedByCoordinate --outSAMunmapped Within --quantMode TranscriptomeSAM --outSAMattributes Standard. Ribosomal RNA features were removed from the GTF file prior to alignment. Aligned BAM files were quantified using featureCounts (v0.46.1)[61], specifying the ribosomal RNA-filtered GTF file to obtain gene-level counts. These counts were then analysed for differential expression using DESeq2 (v1.43.5)[62]. A significance threshold of adjusted p-value ≤ 0.05 and |log₂ fold change| ≥ 0.58 was applied. Principal component analysis (PCA) was performed on variance-stabilised transformed RNA-seq counts generated using DESeq2. Volcano plot and heatmaps were generated using custom R scripts. Gene Ontology (GO) analysis was conducted using ShinyGO (v0.76) [63].

### Bioinformatic analysis of transcription factors using the ISMARA platform

To assess transcription factor (TF) activity from RNA-seq data, we utilized the Integrated System for Motif Activity Response Analysis (ISMARA) platform. For ISMARA analysis, gene expression data were provided as input. The analysis was conducted using the ISMARA web interface, selecting the appropriate genome assembly (Mus musculus GRCm39) and specifying the inclusion of miRNA regulation. ISMARA inferred TF and miRNA activities across samples, identified enriched gene categories among their targets, and predicted direct interactions between regulators (https://ismara.unibas.ch/ISMARA/scratch/data_kgk2a_ti/ismara_report/). The results were visualised using ISMARA’s built-in tools, including motif activity profiles, Z-value bar charts, regulatory motif top targets, and gene category enrichment analyses.

### Dual Luciferase assay

To assess NFAT activity, cells were co-transfected with NFAT-Luc, a firefly luciferase reporter plasmid, and pRLMCV, a Renilla luciferase control, using Lipofectamine™ 3000 Transfection Reagent (Invitrogen, USA) according to the manufacturer’s protocol. After 48 hours, cells were lysed, and Firefly luciferase activity was measured by adding Luciferase Assay Reagent II (Promega, Cat# E1910) to the cell lysate, and luminescence was recorded. Following this, Renilla luciferase activity was measured by adding Stop & Glo® Reagent (Promega, Cat# E1920) to the same sample, and luminescence was recorded again. Data were normalised by calculating the firefly-to-Renilla luciferase activity ratio to account for transfection efficiency.

### Chromatin Immunoprecipitation (ChIP)

ChIP was performed using WT mice and PARP16KO mice’s heart tissue as per the protocol adapted from a previous study [46, 47]. The heart tissues were homogenized and then cross-linked using 1% formaldehyde prepared in 1X PBS for 15 minutes. Formaldehyde was further quenched by using 125mM glycine for 5 minutes at room temperature. Samples were harvested by centrifugation at 2000g for 10 minutes at 4°C. Sonication was performed using Diagenode bioruptor cycles and was fixed at the given setting 25 sec on and 40 sec off for a total of 15 cycles. Chromatin was further visualized on an agarose gel, and it ranged from 200bp to 600bp. For Immunoprecipitation, 50 μg of chromatin was used.

### Molecular modelling

Missing residues in the crystal structures of PARP16 (PDB ID: 6HXS) and NFAT1 (PDB ID: 1A02) were reconstituted using AlphaFold2 [33]. The amino acid sequence of PARP16 (Accession: Q8N5Y8; residues 1-279) and NFAT1 (Accession: Q13469; residues 391-684) were retrieved from UniProt in FASTA format. Structure predictions were performed using a locally installed version of ColabFold (v1.5.2) [48] with GPU acceleration. Multiple sequence alignments (MSAs) were generated with MMseqs2 against the UniRef30 and environmental sequence databases (mmseqs2_uniref_env). Sequences from the same species were paired, and the max-msa parameter was adjusted to 512:1024 to control alignment depth. For template-guided modelling, the PDB100 database was used to retrieve structural homologs. For each protein, predictions were run with 12 recycles (num-recycle = 12), with early stopping enabled when the pLDDT deviation fell to 0 across consecutive recycles (recycle-early-stop-tolerance = 0). To improve conformational sampling, stochastic dropout was activated (use-dropout = True). Each prediction used 20 random seeds (num-seeds = 20), generating five models per seed, yielding a total of 100 models per protein. All models were subjected to AMBER force-field relaxation, comprising 2000 minimisation steps, to reduce steric clashes and optimise stereochemical quality. The top-ranked relaxed models of PARP16 and NFAT1 based on pLDDT and structural quality metrics were selected for downstream analyses.

Blind protein-protein docking of PARP16 against NFAT1 was performed using multiple independent docking platforms: ClusPro [34], HDOCK [35], LightDock [36], and LzerD [37], which generated 120, 100, 100, and 500 docked models, respectively. All models were ranked using their native scoring functions and clustering criteria. Interfacial residues within 8 Å were identified using custom Python scripts. Docking of PARP16 with NAD was performed using the AlphaFold3 server (https://alphafoldserver.com/) [49]. The resulting PARP16-NAD complex was structurally aligned with the top-ranked ClusPro-generated model of PARP16-NFAT1 using the MatchMaker tool in UCSF Chimera [50] to construct the ternary PARP16-NAD-NFAT1 complex for subsequent molecular dynamics simulations.

### All-atom molecular dynamics simulations

Simulation systems were built using the CHARMM-GUI input generator [51]. Solvation was performed with TIP3P water molecules in a cubic box with an edge distance of 89 Å. Counter-ions were added to neutralise the system, and additional NaCl was added to a physiological concentration of 0.15 M. Simulations were carried out using GROMACS (v2023.3)[52] with the CHARMM36m force-field [53]. Energy minimisation was carried out using the steepest-descent algorithm for up to 50,000 steps or until the maximum force converged below 100 kJ mol⁻¹ nm⁻¹. Bond lengths involving hydrogen atoms were constrained using the LINCS algorithm, and the Verlet cutoff scheme was used for non-bonded interactions. Following minimisation, a two-step equilibration protocol was applied. NVT ensemble equilibration was performed for 1 ns at 303.15 K using the V-rescale thermostat. NPT ensemble equilibration was performed for 1 ns at 303.15 K and 1 bar, maintained with the Parrinello-Rahman barostat. After equilibration, all positional restraints were removed, and a 1000 ns production simulation was performed in the NPT ensemble at 303.15 K and 1 bar using a 2 fs integration time step. Long-range electrostatics were calculated using the Particle Mesh Ewald (PME) method. Trajectories were saved every 100 ps, producing 10,000 frames for analysis.

Intermolecular residue-residue contact distances were computed using MDAnalysis (v2.8.0) [54]. Average structures of the ternary complexes were generated using bio3d (v2.4-5) [55]. Non-covalent interactions were characterised with ProLIF (v2.0.3) [56]. MM-PBSA-based binding free energy calculations were carried out using gmx_MMPBSA (v1.6.4) [57]. Three-dimensional structural representations were rendered in UCSF Chimera [50] and figures were prepared using custom R scripts.

### Mass spectrometry analysis

Immunoprecipitated samples were resolved by SDS-PAGE, stained, and excised at the expected molecular weight regions for in-gel digestion with sequencing-grade chymotrypsin. Extracted peptides underwent nano-liquid chromatography-tandem mass spectrometry (nano LC-MS/MS) analysis. Peptides separated on a 110-min gradient (2–90% mobile phase B) at 0.5 μL/min, using mobile phase A (0.1% formic acid in water) and B (0.1% formic acid in 80% acetonitrile). We acquired full MS scans (m/z 350–2000) at 60,000 resolution in positive-ion mode via nano-electrospray on an Orbitrap in data-dependent acquisition mode, with HCD fragmentation of the top 20 precursors and dynamic exclusion enabled. Raw files were searched against UniProt (chymotrypsin enzyme; ADP-ribosylation as variable modification), filtering peptide identifications to <1% FDR.

### Statistical analysis

All data are presented as mean ± SD unless otherwise indicated. Data distribution was assessed for normality using the Shapiro-Wilk test. For comparisons between 2 groups, a 2-tailed unpaired Welch’s t test was used for normally distributed data with unequal variances. For non-normally distributed data, the Mann-Whitney U test was applied. For comparisons among 3 or more groups, 1-way ANOVA followed by a Tukey multiple-comparisons test. For experiments involving 2 independent variables, 2-way ANOVA followed by Sidak multiple-comparisons test was used. Normality for ANOVA analyses was assessed within each group. The value of n represents independent biological replicates (individual animals or independently prepared primary cardiomyocyte cultures). Technical replicates were averaged and treated as a single biological replicate. A p-value of <0.05 was considered statistically significant. Statistical analyses were performed using GraphPad Prism version 8.4.3.

---

**Figure S1.**
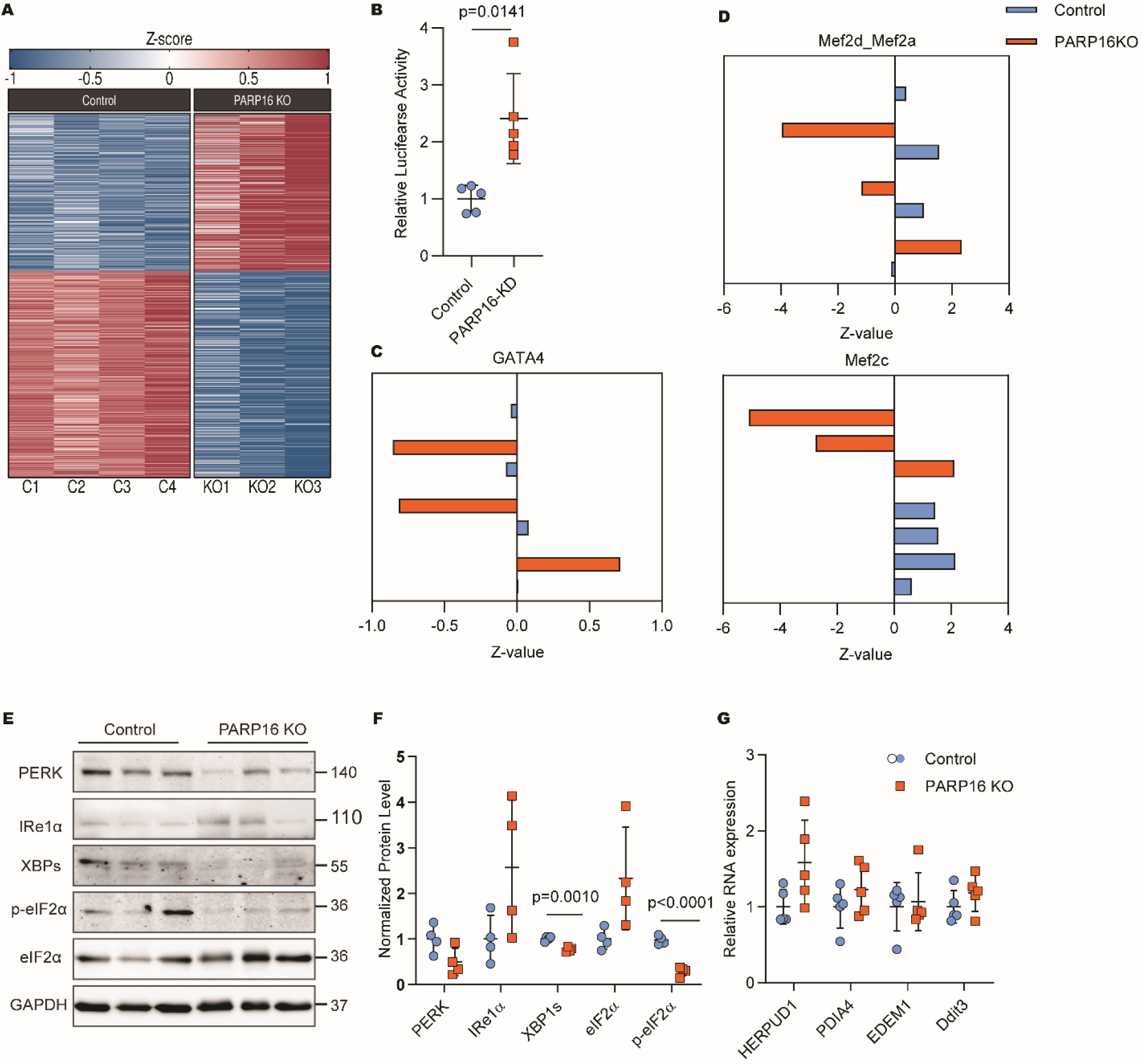
Additional transcriptomic analyses and unfolded protein response signalling in control and PARP16-deficient hearts. (A) Heatmap showing all differentially expressed genes (DEGs) identified by RNA sequencing analysis of heart tissue from 5-month-old control and PARP16 knockout (KO) mice. (B) NFAT1 luciferase reporter activity in neonatal rat cardiomyocytes (NRCMs) transfected with control siRNA or PARP16-targeting siRNA. (C) ISMARA-predicted GATA4 transcription factor activity in individual 5-month-old control and PARP16 KO heart samples derived from RNA sequencing data. (D) ISMARA-predicted MEF2 family transcription factor activity (MEF2D/MEF2A and MEF2C) in individual 5-month-old control and PARP16 KO heart samples derived from RNA sequencing data. (E) Representative immunoblot analysis of unfolded protein response (UPR) signalling proteins in hearts from 5-month-old control and PARP16 KO mice. Molecular weight markers are indicated. (F) Quantification of UPR-associated protein expression from independent experiments (n = 4 mice per group). (G) Quantitative RT-PCR analysis of UPR target gene expression in hearts from 5-month-old control and PARP16 KO mice (n = 5 mice per group). Data are presented as mean ± SD. Statistical analyses were performed as indicated in the figure. Exact P values are provided in the source data. Source data are provided as a Source Data file.

**Figure S2.**
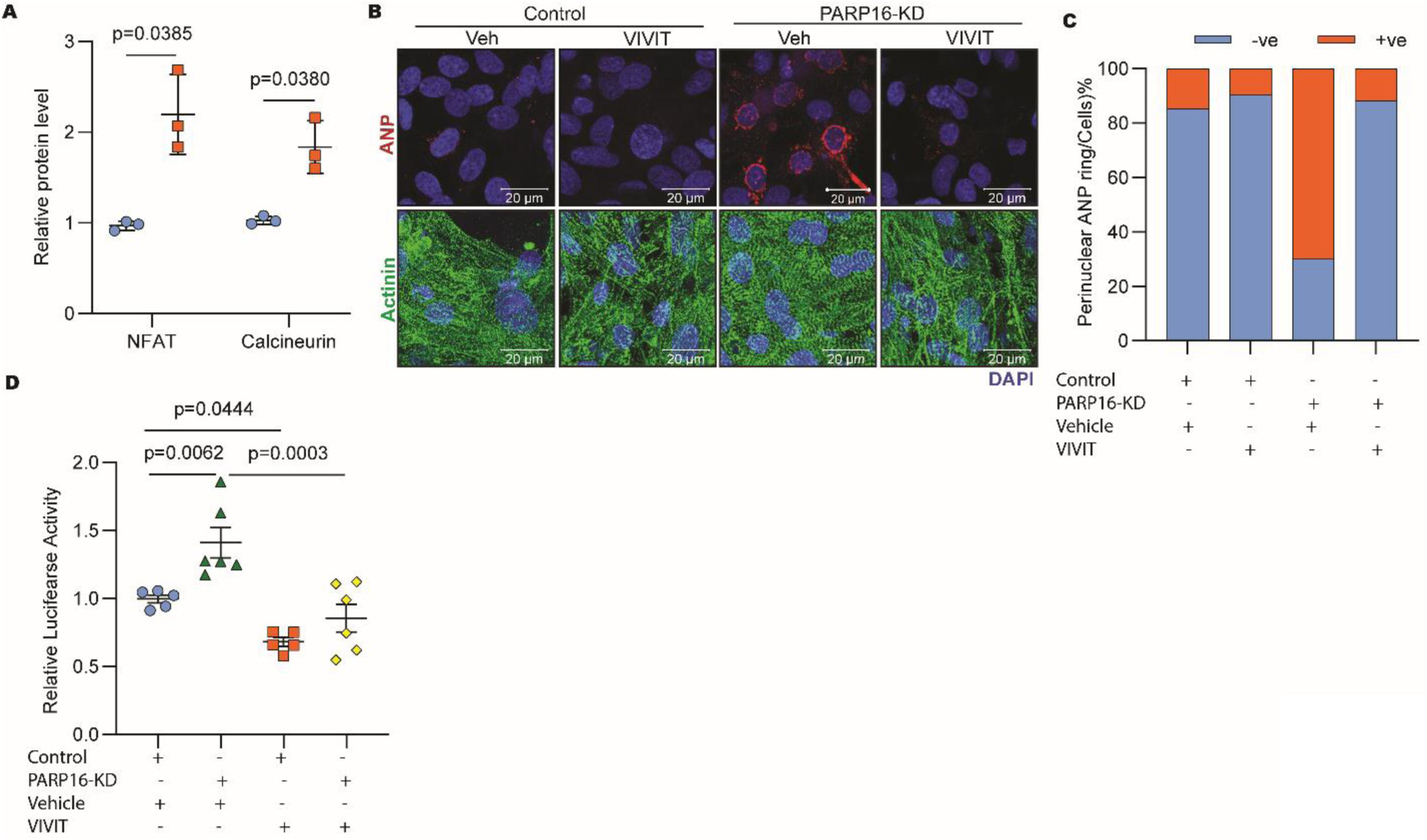
Modulation of NFAT1 activity and cardiomyocyte remodelling by pharmacological inhibition of NFAT signalling. (A) Quantification of total NFAT1 and calcineurin protein levels in hearts from 5-month-old control and PARP16 knockout (KO) mice. Protein abundance was normalised to GAPDH (n = 3 mice per group). (B) Representative confocal images of neonatal rat cardiomyocytes (NRCMs) transfected with control siRNA or PARP16-targeting siRNA and treated with vehicle or the NFAT activator inhibitor VIVIT. Cells were immunostained for atrial natriuretic peptide (ANP), α-actinin, and DAPI. Scale bar, as indicated in the figure. (C) Quantification of the percentage of NRCMs exhibiting perinuclear ANP localisation under control or PARP16 knockdown conditions following vehicle or VIVIT treatment. (D) NFAT1 luciferase reporter activity in control and PARP16-knockdown NRCMs treated with vehicle or VIVIT (n = 6 independent experiments). Data are presented as mean ± SD. Statistical analyses were performed as indicated in the figure. Exact P values are provided in the source data. Source data are provided as a Source Data file.

**Figure S3.**
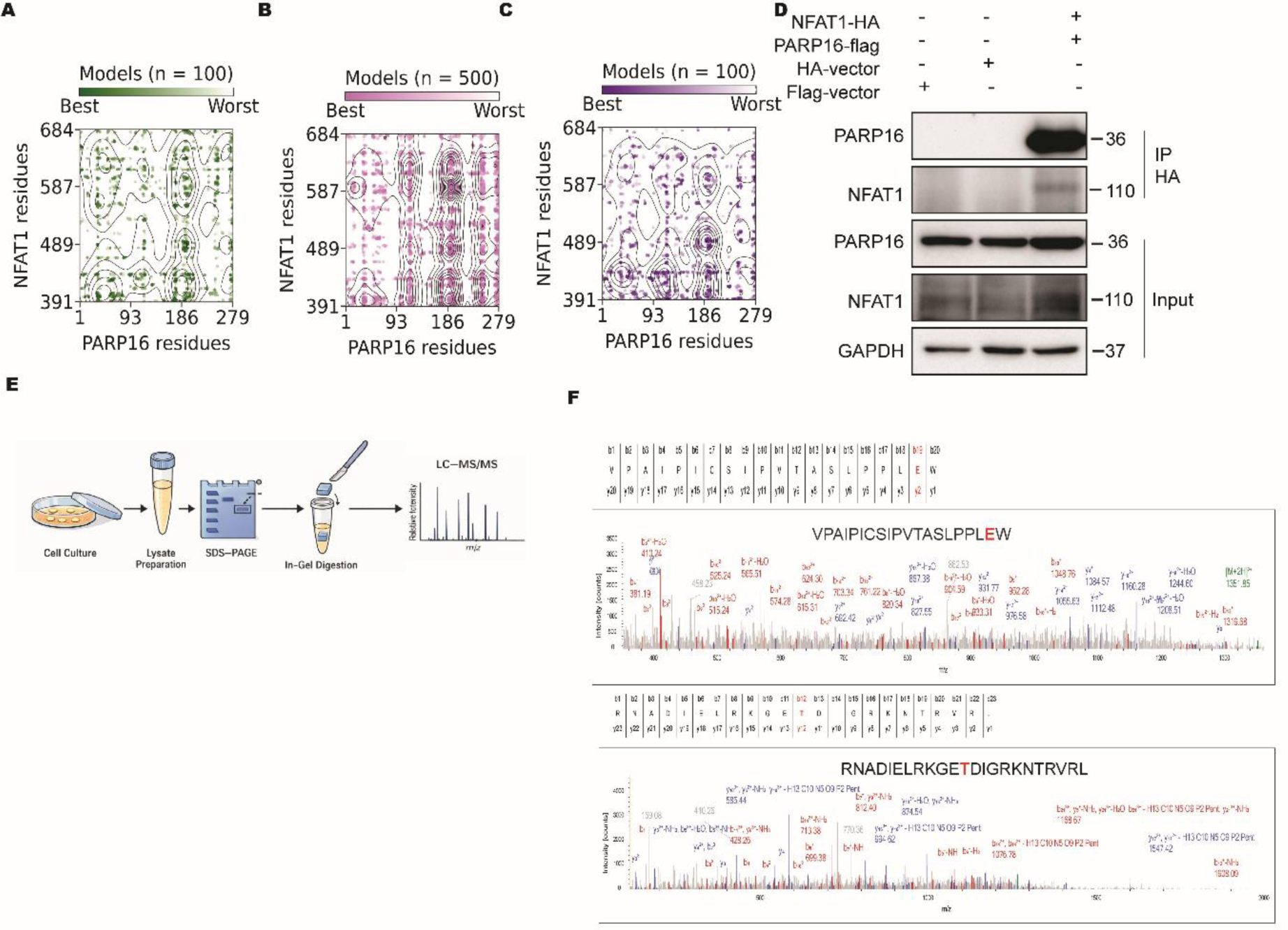
Supporting analyses of PARP16–NFAT1 interaction and identification of candidate NFAT1 residues. (A) Distribution of interfacial PARP16–NFAT1 residue pairs identified within 8 Å across 100 HDOCK-generated docking models. Point colour distinguishes residue pairs derived from top-ranked and lower-ranked docking solutions. (B) Distribution of interfacial PARP16–NFAT1 residue pairs identified within 8 Å across 500 LZerD-generated docking models. Point colour distinguishes residue pairs derived from top-ranked and lower-ranked docking solutions. (C) Distribution of interfacial PARP16–NFAT1 residue pairs identified within 8 Å across 100 LightDock-generated docking models. Point colour distinguishes residue pairs derived from top-ranked and lower-ranked docking solutions. (D) Immunoblot analysis of haemagglutinin (HA) immunoprecipitation in HEK293 cells expressing Flag vector, HA vector, or PARP16-FLAG together with HA-NFAT1. Immunoprecipitated complexes were analysed by immunoblotting with the indicated antibodies. (E) Schematic illustration of the mass spectrometry workflow used for identification of candidate NFAT1 residues following PARP16 overexpression. (F) Representative mass spectrometry spectra corresponding to NFAT1 peptides containing candidate residues E398 (top) and T533 (bottom) identified following PARP16 WT overexpression. Data are provided as a Source Data file.

